# Phosphorylation-dependent mitotic SUMOylation drives nuclear envelope-chromatin interactions

**DOI:** 10.1101/2021.03.24.436817

**Authors:** Christopher Ptak, Natasha O. Saik, Ashwini Premashankar, Diego L. Lapetina, John D. Aitchison, Ben Montpetit, Richard W. Wozniak

**Author notes:** These authors contributed equally (Christopher Ptak, Natasha O. Saik).

## Abstract

In eukaryotes, chromatin binding to the inner nuclear membrane (INM) and nuclear pore complexes (NPCs) contributes to spatial organization of the genome and epigenetic programs important for gene expression. In mitosis, chromatin-nuclear envelope (NE) interactions are lost and then formed again as sister chromosomes segregate to post-mitotic nuclei. Investigating these processes in *S. cerevisiae*, we identified temporally and spatially controlled phosphorylation-dependent SUMOylation events that positively regulate post-metaphase chromatin association with the NE. Our work establishes a phosphorylation-mediated targeting mechanism of the SUMO ligase Siz2 to the INM during anaphase, where Siz2 binds to and SUMOylates the VAP protein Scs2. The recruitment of Siz2 through Scs2 is further responsible for a wave of SUMOylation along the INM that supports the assembly and anchorage of subtelomeric chromatin at the INM and localization of an active gene (*INO1*) to NPCs during the later stages of mitosis and into G1-phase.

## Introduction

The spatial organization of a eukaryotic genome within the nucleoplasm is influenced by multiple factors including the nuclear envelope (NE), which functions as a 2-dimensional interaction surface for chromatin. The inner nuclear membrane (INM) often associates with densely packed, gene poor, and transcriptionally inactive heterochromatin, while nuclear pore complexes (NPCs) typically engage transcriptionally active euchromatin (Ptak et al., 2014; Buchwalter et al., 2019; Misteli, 2020). However, these interactions are not static, and in multiple eukaryotes it has been shown that they are remodeled in response to gene activation, DNA metabolism (e.g., replication or repair), and during mitosis (e.g., NE breakdown and chromosome segregation) (Güttinger et al., 2009; Taddei and Gasser, 2012; Ptak et al., 2014; Ma et al., 2015; Ptak and Wozniak, 2016). In *S. cerevisiae*, heterochromatin-like subtelomeric chromatin is associated with the INM and transcriptional activation has been shown to involve localization of specific gene loci to NPCs (e.g., *INO1* and *GAL1-10*) (Egecioglu and Brickner, 2011; Taddei and Gasser, 2012). In addition, although the NE remains intact during mitosis in budding yeast, chromatin-NE interactions are also lost and reformed during mitosis (Hediger et al., 2002; Ebrahimi and Donaldson, 2008; Brickner and Brickner, 2010). This overall regulation of chromatin-NE interactions, i.e., both disruption and reformation of these contacts, is critical for proper genome organization and epigenetic inheritance during cell division (Champion et al., 2019; Falk et al., 2019; Poleshko et al., 2019; Politz et la., 2015).

As cell progress through mitosis, the cyclical phosphorylation/dephosphorylation of various NE proteins contributes to the disruption and later reformation of NE-chromatin interactions (Güttinger et al., 2009; Wurzenberger and Gerlich, 2011). Despite the importance of re-establishing chromatin-NE interactions in the later stages of mitosis, it is unclear whether these events are supported by specific processes, such as phosphorylation or other post-translational modifications (PTMs). A candidate for such a functional role is SUMO (small ubiquitin-like modifier) modification. SUMOylation has been linked to heterochromatin assembly and chromatin tethering in yeast and higher eukaryotes (Hari et al., 2001; Ferreira et al., 2011; Lapetina et al., 2017; Ninova et al., 2019) and the association of active genes with NPCs (Texari and Stutz, 2015; Saik et al., 2020). In addition, the SUMO E3 ligase Siz2, has been implicated in the formation of NE-chromatin interactions during interphase (Ferreira et al., 2011; Freudenreich and Su, 2016; Lapetina et al., 2017; Saik et al., 2020).

Here, we report a PTM cascade involving phosphorylation and SUMOylation that is essential for establishing NE-chromatin interactions in the later stages of mitosis. With anaphase onset, we find that phosphorylation induces binding of Siz2 to the VAP (<u>V</u>esicle-associated membrane protein (VAMP)-<u>A</u>ssociated <u>P</u>rotein) family member Scs2 at the INM where it remains until cytokinesis. At the INM, Siz2 drives a wave of mitotic SUMOylation involving NE proteins, with targets that include Scs2 and subtelomeric chromatin associated Sir4. We show that these spatially regulated SUMOylation events are required to establish the proper organization of telomeres at the INM and the association of the activated *INO1* gene with NPCs by anaphase/telophase of mitosis. These findings establish a temporally and spatially regulated set of phosphorylation-dependent SUMOylation events that regulate the re-establishment of nuclear architecture during the late stages of mitosis.

## Results

### Scs2 directs mitotic SUMOylation at the NE

Immunofluorescence (IF) microscopy and western blotting were performed using a SUMO-specific antibody to investigate spatial and temporal changes in cellular SUMOylation during the cell cycle in *S. cerevisiae*. Imaging of asynchronous IF stained cells revealed a predominantly nuclear SUMO signal in interphase cells (unbudded and small-budded cells) and a concentration of SUMO at septin rings in mitotic cells (large budded) as previously reported (Johnson and Gupta, 2004). However, in mitotic cells, SUMO conjugates were also concentrated at the NE (Fig. 1 A). Coinciding with this mitotic redistribution of SUMO, western blotting analysis in synchronized cell cultures revealed an increase in mitotic levels of various SUMOylated species that occurred just after peak levels of the cyclin Clb2 (Fig. 1 B). Noticeable among these were four clearly resolved SUMOylated species in the 40-55 kDa range, which are smaller in size than the septins that are known to be SUMOylated in mitosis (Johnson and Gupta, 2004). Given that Clb2 is degraded upon anaphase onset (Irniger, 2002), the combined data are suggestive of SUMOylation events occurring in late metaphase/anaphase when SUMO is evident at the NE.

**Figure 1.**
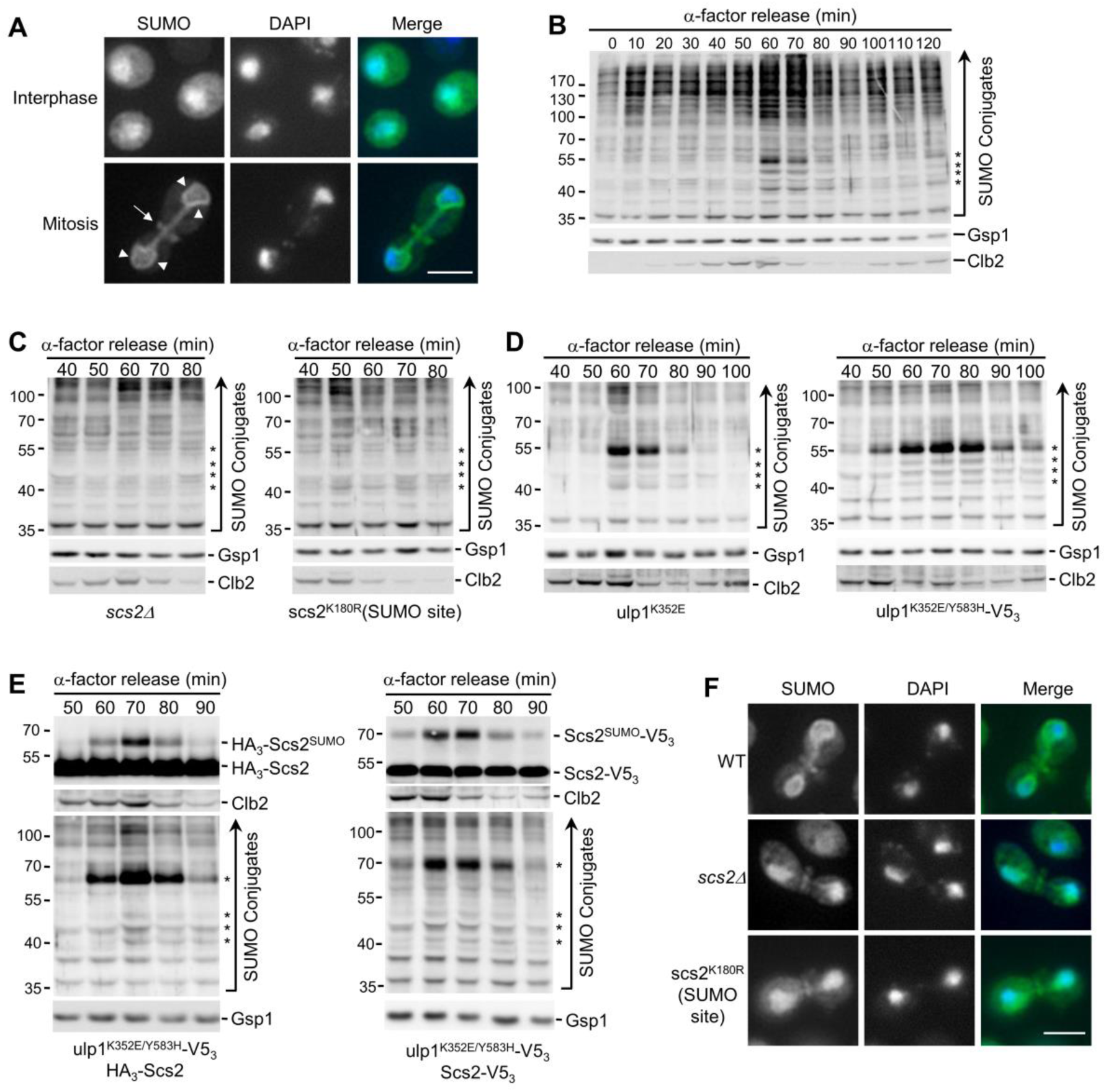
Scs2 directs mitotic SUMOylation events. (**A** and **F**) Epifluorescence images of WT (**A, F**), *scs2Δ*, and *scs2^K180R^* (SUMOylation site mutant - SUMO site) (**F**) cells analyzed by anti-SUMO immunofluorescence (SUMO). Nuclear position is determined by DAPI staining. Arrowheads highlight SUMOylated proteins along the NE. SUMOylated septin ring position is indicated by an arrow. Images were rendered using the unsharp mask filter in Image J. Bar, 2μm. (**B – E**) Cells were arrested in G1-phase using α-factor. Following α-factor removal, cultures were sampled every 10 min and analyzed by western blotting using antibodies directed against the proteins indicated on the right. Note, Clb2 levels peak in metaphase. All strains used are *bar1Δ* and the presence of a mutation and/or tagged protein is indicated. Gsp1 is a loading control. Asterisks identify the expected position of mitotic, Scs2-dependent SUMO conjugates. Molecular mass markers are shown in kDa.

To identify these prominent mitotic targets, we curated a list of SUMOylated proteins of similar masses (Panse et al., 2004; Wohlschlegel et al., 2004; Wykoff and O’Shea, 2005; Zhou et al., 2004; Denison et al, 2005; Hannich et al., 2005). From this list of proteins, 141 nonessential genes were selected, and an analysis of null mutant strains revealed that the four mitotic SUMOylated species were absent in cells lacking the gene encoding Scs2 (Fig. 1 C). Scs2 is an ER/NE localized membrane protein of the VAP (<u>V</u>esicle-associated membrane protein (VAMP)- <u>A</u>ssociated <u>P</u>rotein) family (Loewen and Levine, 2005). Consistent with this observation, mutation of a known SUMO acceptor site in Scs2 (Felberbaum et al., 2012), scs2^K180R^, also eliminated, or reduced the level of, these SUMOylated species (Fig. 1 C).

Two further approaches were used to confirm the identity of Scs2 as a mitotic SUMOylation target. First, we employed a mutation in the deSUMOylase Ulp1 (*ulp1^K352E^*) shown to increase cellular levels of SUMOylated Scs2 (Felberbaum et al., 2012). As seen in Fig. 1 D, the *ulp1^K352E^* mutant showed a marked increase in the level of the ∼55 kDa SUMOylated protein during mitosis. Serendipitously, we discovered a V5_3_-tagged *ULP1* allele bearing an additional mutation (ulp1^K352E/Y583H^-V5_3_) that showed further elevated levels of Scs2-SUMO (Fig. 1 D). Second, N- or C-terminal tagging of endogenous Scs2 specifically shifted the electrophoretic mobility of the ∼55 kDa SUMOylated species (Fig. 1 E). These results suggest that the three smaller SUMOylated species migrating faster than the ∼55 kDa species are not proteolytic fragments of Scs2-SUMO and likely represent other mitotic SUMOylation targets that are dependent on Scs2 SUMOylation. Consistent with this conclusion, we also observed that upon loss of Scs2, or in strains containing the scs2^K180R^ SUMO site mutant, SUMO accumulation at the NE during mitosis was not observed (Fig. 1 F). We conclude that Scs2 is a NE-associated mitotic SUMOylation target that directs mitotic SUMOylation events in a manner dependent upon its own SUMOylation.

### Siz2 recruitment to the NE directs mitotic SUMOylation

In *S. cerevisiae* three SUMO E3 ligases, Siz1, Siz2/Nfi1, and Mms21 (Jentsch and Psakhye, 2013), guide the selection of most SUMOylation targets in actively growing cells. Anti-SUMO western blotting of synchronized or asynchronous cell cultures showed that only cells lacking Siz2 failed to accumulate Scs2-SUMO and smaller, prominent SUMOylated species during mitosis (Fig. 2 A and Fig. S1 A, 1 B). Moreover, Siz2, but not Siz1 or Mms21, was required for mitotic NE accumulation of SUMO conjugates (Fig. 2 B, Fig. S1 C). Together, these observations indicate that Siz2, as well as Scs2, are required to direct mitotic SUMOylation events at the NE.

**Figure 2.**
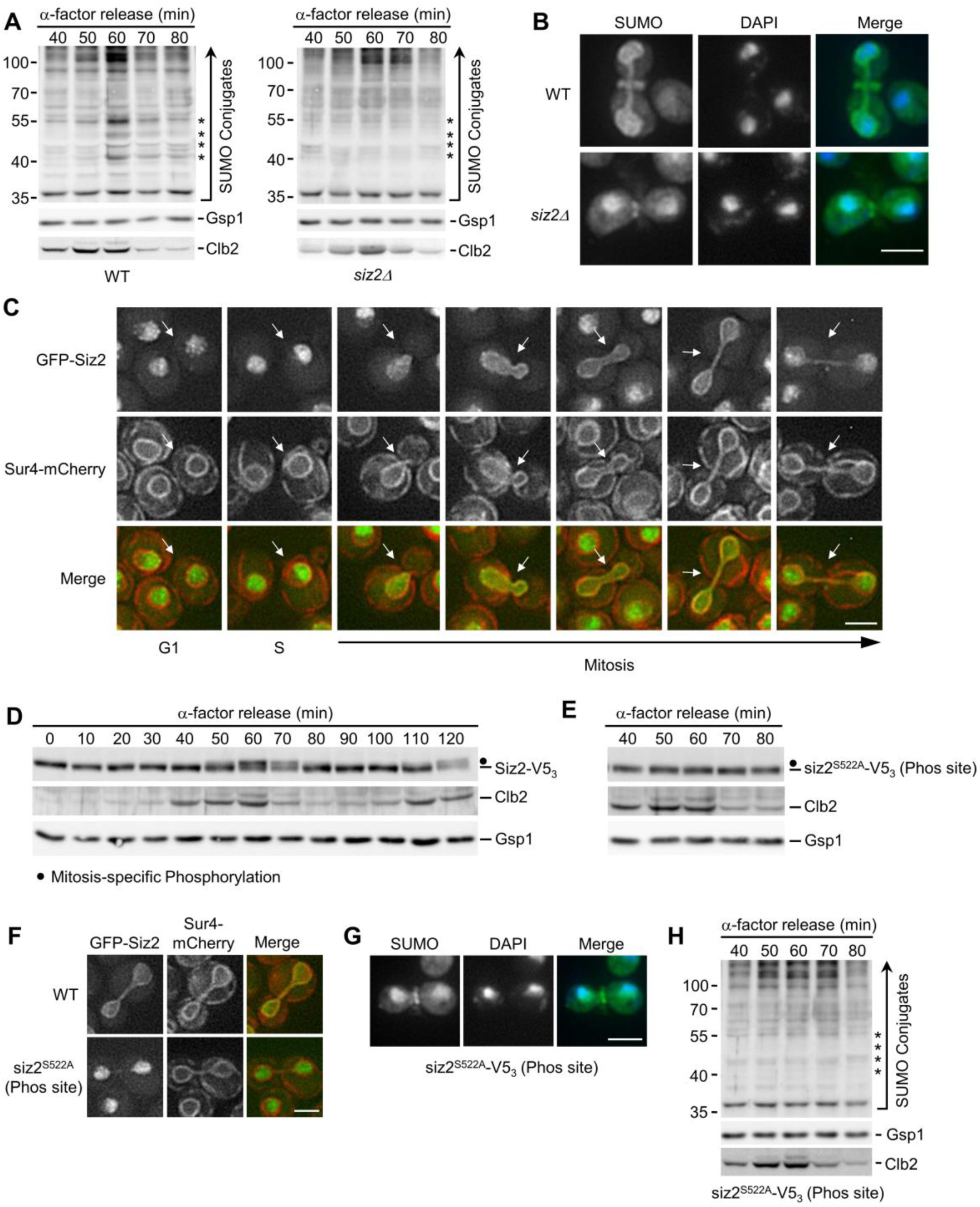
Mitotic recruitment of Siz2 to the NE directs SUMOylation events. (**A**, **D**, **E**, and **H**) α-factor arrest-release assays were carried out as in Fig. 1. Cell lysates were analyzed by western blotting to detect SUMO conjugates (**A**, **H**), Siz2-V5_3_ (**D**) and siz2^S522A^-V5_3_ (phosphorylation site mutant - Phos site) (**E**), as well as Clb2, and the Gsp1 load control in the indicated strain background. Relevant information on each mutant is provided. Asterisks identify the expected position of mitotic, Scs2-Siz2 dependent SUMO conjugates. Dots indicate the expected position of mitotically phosphorylated Siz2-V5_3_. Molecular mass markers are shown in kDa. (**B** and **G**) Anti-SUMO immunofluorescence analysis of WT and *siz2Δ* (**B**) as well as *siz2^S522A^* (**G**) cells was carried out as described in Fig. 1. DAPI staining identifies nuclear position. (**C** and **F**), Epifluorescence images of cells producing GFP-Siz2 (**C**, **F**) or GFP-siz2^S522A^ (**F**) along with the NE/ER marker Sur4-mCherry. Images were rendered using the unsharp mask filter in Image J. Cell cycle stage of each highlighted cell (arrows) is indicated in (**C**). Mitotic cells are shown in (**F**). Bar, 2μm.

Siz1 has previously been shown to target to the septin ring during M-phase where it SUMOylates the septins (Johnson and Gupta, 2001) (see Fig. S1). In an analogous manner, we hypothesized that the spatiotemporal recruitment of Siz2 to the NE during mitosis may direct SUMOylation of NE proteins. Upon examination of GFP-Siz2 localization in G1- and S-phase cells, we observed GFP-Siz2 diffusely, and in distinct puncta, throughout the nucleoplasm, while partially excluded from the nucleolus (Fig. 2 C, Fig. S1 D). GFP-Siz2 puncta were also visible along the NE, including the NE adjacent to the nucleolus. Time-lapse imaging revealed that these puncta are dynamic and do not persist in mitosis (Video 1). Strikingly, as the NE elongated and cells progressed into anaphase, GFP-Siz2 accumulated at the NE where it remained until dissolution of the NE membrane bridge that links the mother and daughter nuclei during cytokinesis (Fig. 2 C, Fig. S1 E, and Video 1).

Critical transitions throughout mitosis are driven by PTMs, including phosphorylation (Cuijpers and Vertegaal, 2018). Given the sharp transition of GFP-Siz2 from the nucleoplasm to the NE, we examined whether Siz2 was post-translationally modified during mitosis using western blot analysis of synchronized cells (Fig. 2 D). Between 60- and 70-minutes post G1-phase arrest, we observed a change in the electrophoretic mobility of Siz2 (tagged with V5_3_ for detection) consistent with it being post-translationally modified. Notably, this mobility change corresponded with the peak mitotic levels and subsequent loss of Clb2, suggesting that Siz2-V5_3_ was modified late in metaphase or at the onset of anaphase (Irniger, 2002). These data indicate that a Siz2 PTM occurs at the same time as Siz2 NE recruitment and Scs2 SUMOylation (Fig. 1 and 2).

Phosphoproteome analyses have previously identified putative Siz2 phosphorylation sites at serine residues 522, 527, and 674 (Albuquerque et al., 2008; Holt et al., 2009). Consistent with Siz2 being phosphorylated, phosphatase treatment of mitotic cell lysates removed the slower migrating species (Fig. S1 F). By examining Siz2 electrophoretic mobility in serine to alanine point mutants, siz2^S527A^-V5_3_ and siz2^S674A^-V5_3_ mutants were found to retain the electrophoretic shift during mitosis similar to Siz2-V5_3_ (Fig. S1 G), but this shift was absent from the siz2^S522A^- V5_3_ mutant (Fig. 2 E). Importantly, GFP-siz2^S522A^ failed to localize to the NE during mitosis (Fig. 2 F), *siz2^S522A^* mutant cells did not accumulate SUMO at the NE in mitosis (Fig. 2 G), and anti-SUMO western blotting analysis showed loss of the four prominent mitotic SUMOylated species in *siz2^S522A^* including Scs2 (Fig. 2 H). By contrast, *siz2^S527A^* and *siz2^S674A^* mutants showed SUMO-conjugate profiles similar to WT cells (Fig. S1 G). These results suggest that phosphorylation-dependent recruitment of Siz2 to the NE leads to SUMOylation of Scs2 and several other NE proteins during mitosis.

### Scs2 is required for Siz2 recruitment to the NE

The shared phenotypes of Scs2 and Siz2 mutants suggested to us that they may form a complex at the NE to direct mitotic SUMOylation events. Although Scs2 functions as a receptor on the ER for multiple cytoplasmic proteins (Stefan et al., 2011; Chao et al, 2014; Encinar Del Dedo et al., 2017; Ng et al, 2020), our postulate would require that Scs2 is present at the INM to bind nucleoplasmic Siz2. Scs2 contains a single transmembrane region at its C-terminus that tethers it to the ER and ONM (Loewen and Levine, 2005), but an INM pool of Scs2 has not been described. To assess Scs2 localization, we used a split superfolder GFP assay previously used to characterize the INM proteome (Smoyer et al., 2016). In this assay, if a GFP_11_-reporter resides in the same subcellular compartment as a target protein fused to GFP_1-10_, the two GFP fragments can assemble and fluoresce. As expected, we observed a pool of Scs2 localized at the ER/ONM, based on the association of GFP_1-10_-Scs2 with the cytoplasmic GFP_11_-Hxk1 reporter (Fig. 3 A). Importantly, GFP_1-10_-Scs2 and the nucleoplasmic GFP_11_-Pus1 reporter also yielded a robust NE GFP signal, but no ER signal, suggesting Scs2 accesses the INM (Fig. 3 A). Consistent with Scs2 acting as a receptor for Siz2 at the INM, Siz2 was phosphorylated but no longer localized to the NE during mitosis in cells lacking Scs2 (*scs2Δ*) or producing a scs2^1-225^ truncation missing the transmembrane domain (Brickner and Walter, 2004; Loewen et al., 2007) (Fig. 3 C and Fig. S2 A). Furthermore, immunoprecipitation followed by western blotting showed that Scs2-TAP binds Siz2-V5_3_, but only weakly to siz2^S522A^-V5_3_ (Fig. 3 B). These observations support the conclusion that Scs2 is an INM receptor for phosphorylated Siz2 during late stages of mitosis.

**Figure 3.**
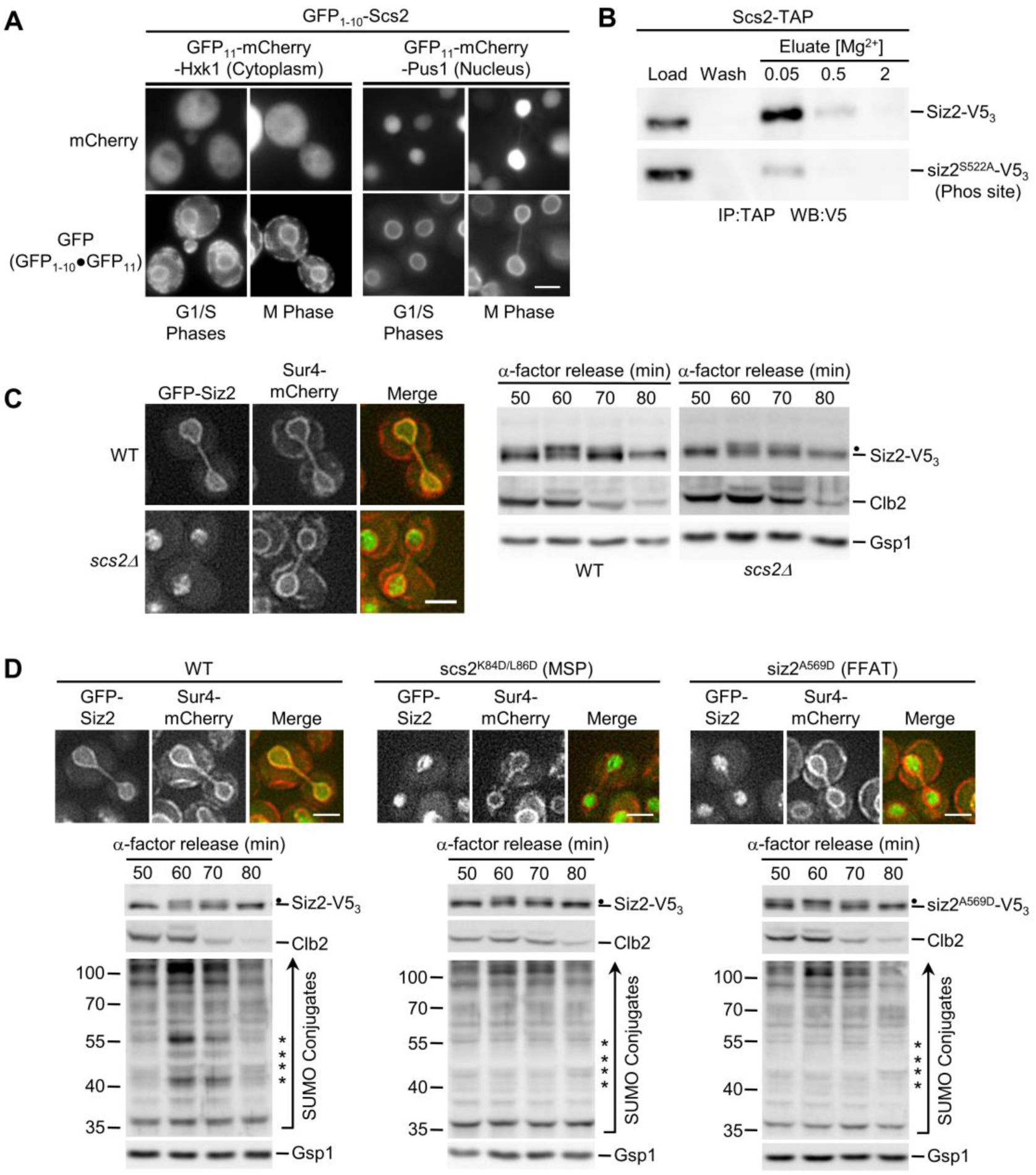
INM association of Scs2-Siz2 directs mitosis specific SUMOylation events. (**A**) Scs2 localization was assessed using the split-superfolder GFP system. Epifluorescence images of WT cells containing GFP_1-10_-Scs2 and plasmid-encoded GFP_11_-mCherry-Hxk1(cytoplasmic) or GFP_11_-mCherry-Pus1 (nuclear). Localization of the reporter (mCherry) and assembled GFP_1- 10_**·**GFP_11_ (GFP) in representative G1-, S-, and M-phase cells are shown. (**B**) Binding of Siz2-V5_3_ and siz2^S522A^-V5_3_ to Scs2-TAP was assessed. Scs2-TAP was affinity-purified from cells (IP) producing Siz2 derivatives and bound proteins were eluted using a Mg^2+^ step gradient. Indicated fractions were analyzed by western blotting (WB) using an anti-V5 antibody. (**C** and **D)** The indicated strains were assessed for Siz2-V5_3_ phosphorylation and their mitotic SUMO conjugate profile by western blot analysis as described in Figs. 1 and 2. GFP-Siz2 and Sur4-mCherry localization was performed as in Fig. 2. The *scs2^K84D/L86D^* mutations lie in the MSP domain mutant (MSP) and the *siz2^A569^* mutation within the FFAT-like motif (FFAT). Bar, 2μm. Dots indicate the expected position of mitotically phosphorylated Siz2-V5_3_. Asterisks identify the expected position of mitotic, Scs2-Siz2 dependent SUMO conjugates. Molecular mass markers are shown in kDa.

Scs2, like other members of the VAP family, contains a N-terminal MSP (major sperm protein) domain that binds FFAT (two phenylalanines in an acidic tract) or FFAT-like motifs within binding partners (Loewen et al., 2003; Loewen and Levine, 2005; Kaiser et al., 2005). Notably, Siz2 contains three potential FFAT-like motifs (Murphy and Levine, 2016) (Fig. S2 B). Previous studies showed that K84D and L86D substitutions within the MSP motif of Scs2 specifically disrupts interactions with an FFAT-containing peptide (Kaiser et al., 2005). Since Scs2^K84D/L86D^ retains INM localization (Fig. S2 C), we tested the impact of a *scs2^K84D/L86D^* mutant on the NE localization of Siz2. To complement this approach, we also produced a mutation in Siz2 (A569D) predicted to disrupt a putative FFAT-like motif detected between residues 565-571 of Siz2 (Murphy and Levine, 2016). Both the *scs2^K84D/L86D^* and *siz2^A569D^* mutations inhibited mitotic-specific INM localization of Siz2, while not altering mitotic phosphorylation of Siz2 or siz2^A569D^ (Fig. 3 D). The *scs2^K84D/L86D^* and *siz2^A569D^* mutations inhibited SUMOylation of mitotic targets, including Scs2, in both an otherwise WT background (Fig. 3 D) and in *ulp1^K352E/Y583H^-V5_3_* cells (Fig. S2 D). On the basis of these data, we propose that an FFAT-like motif in Siz2 promotes an interaction with the MSP domain of Scs2 at the INM that is critical for Siz2-directed mitotic SUMOylation events.

### A SUMO-interaction motif in Siz2 and SUMOylation of Scs2 contribute to Scs2-Siz2 association

Our accumulated data show that both Siz2 phosphorylation and an FFAT-MSP domain interaction are required for Siz2 to bind and SUMOylate Scs2. This raised the possibility that Scs2 SUMOylation may stabilize its association with Siz2. To investigate this possibility, a strain producing the SUMOylation deficient scs2^K180R^ mutant, which retains INM localization (Fig. S2 E), was examined. In the *scs2^K180R^* mutant, Siz2 was still phosphorylated, but recruitment to the NE was reduced (Fig. 4 A). Given that Siz2 contains two SUMO-interaction motifs (SIM domains), this result suggested that stable binding of Siz2 to Scs2 is supported by a SUMO-SIM interaction. We tested this possibility by examining SIM1 (*siz2^I472/473A^*) or SIM2 (*siz2^V720/721A^*) mutants that would be expected to inhibit SIM function (Psakhye and Jentsch, 2012). Neither mutation changed Siz2 mitotic phosphorylation; however, the SIM1 mutant caused a visible reduction in mitotic NE association of Siz2 and mitotic SUMOylation of Scs2-Siz2 targets, while the SIM2 mutant had little effect (Fig. 4 B). Of note, the mitotic SUMOylation defect of the SIM1 mutant, but not Siz2 localization, was rescued in a strain also producing deSUMOylase defective ulp1^K352E/Y583H^-V5_3_, which is consistent with weak NE association and not a loss of Siz2 SUMO ligase activity (Fig. 4 C). These data demonstrate that the SIM1 motif of Siz2 contributes to Siz2 NE localization and Siz2-mediated SUMOylation during mitosis.

**Figure 4.**
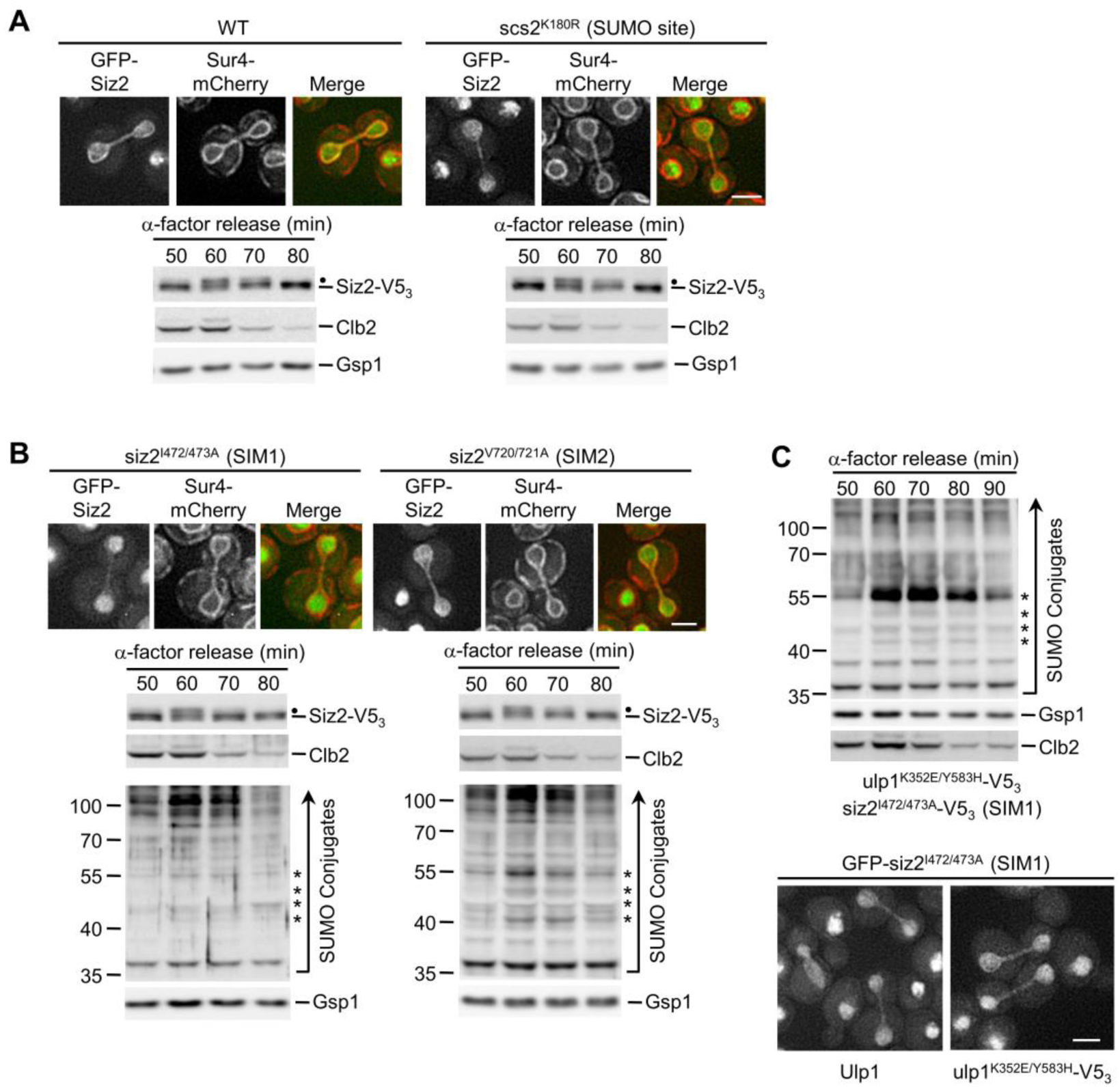
A Siz2 SIM motif and Scs2-SUMO contribute to mitosis-specific SUMOylation. (**A** and **B**) The indicated strains, including WT and mutations in the Scs2 SUMO site (*scs2^K180R^*) or Siz2 SIMs (*siz2^I472/473A^* (SIM1) and *siz2^V720/721A^* (SIM2) were assessed for mitotic Siz2 phosphorylation and GFP-Siz2 and Sur4-mCherry localization as described in Fig. 2. (**B** and **C**) Mitotic SUMO conjugate profiles were assessed as described in Fig. 1 using *siz2^I472/473A^*, *siz2^V720/721A^* (**B**) and *ulp1^K352E/Y583H^-V5_3_ siz2^I472/473A^* (**C**) strains. (**C**) GFP-siz2^I472/473A^ localization in an otherwise WT (Ulp1) or an *ulp1^K352E/Y583H^-V5_3_* strain was assessed. Bar, 2μm. Dots indicate the expected position of mitotically phosphorylated Siz2-V5_3_. Asterisks identify the position of mitotic, Scs2-Siz2 dependent SUMO conjugates. Molecular mass markers are shown in kDa.

### The INM localized Scs2-Siz2 complex is required for telomere tethering to the NE during mitosis

The Scs2-dependent repositioning of Siz2 from the nucleoplasm to the INM is initiated during a period of transition from metaphase to anaphase. During this time, sister chromosomes are segregated to daughter nuclei and specific chromatin-NE interactions are re-established as cells continue into G1-phase of the cell cycle (Hediger et al., 2002; Ebrahimi and Donaldson, 2008). To test for a role of Scs2-Siz2 mediated INM SUMOylation in establishing NE-chromatin interactions during mitosis, we examined the association of telomeres with the INM using a GFP-labeled telomere localization assay (Hediger et al., 2002). In WT cells, telomeres were detected at the NE in ∼65-75% of cells in each of the three cell cycle stages examined: anaphase/telophase, G1-phase, and S-phase. However, *siz2* and *scs2* point mutants that inhibit both Siz2 INM association and SUMOylation of INM targets showed decreased telomere 14L (Tel14L; Fig. 5 A) and 6R (Fig. S3 A) tethering to the NE during anaphase/telophase. Reduced Tel14L tethering persisted into G1- phase, with the exception of the *siz2^A569D^* (FFAT motif) mutation, which showed less of a defect (Fig. 5 A). By contrast, in the *ulp1^K352E/Y583H^-V5_3_* mutant, where Scs2-SUMO levels are elevated in mitosis and into G1-phase (Fig. 1 D), Tel14L levels at the NE are normal in M-phase and increased in G1-phase cells (Fig. S3 B). By S-phase, the NE association of Tel14L in the *siz2*, *scs2*, and *ulp1* point mutants was largely normal or, in the case of *scs2^K180R^* (SUMO site mutation), their levels at the NE increased in S-phase as compared to G1-phase (Fig. 5 A and Fig. S3 B). These data establish a strong correlation between Siz2 localization at the NE, Scs2-Siz2 mediated SUMOylation of INM proteins, and the tethering of Tel14L to the NE during mitosis.

**Figure 5.**
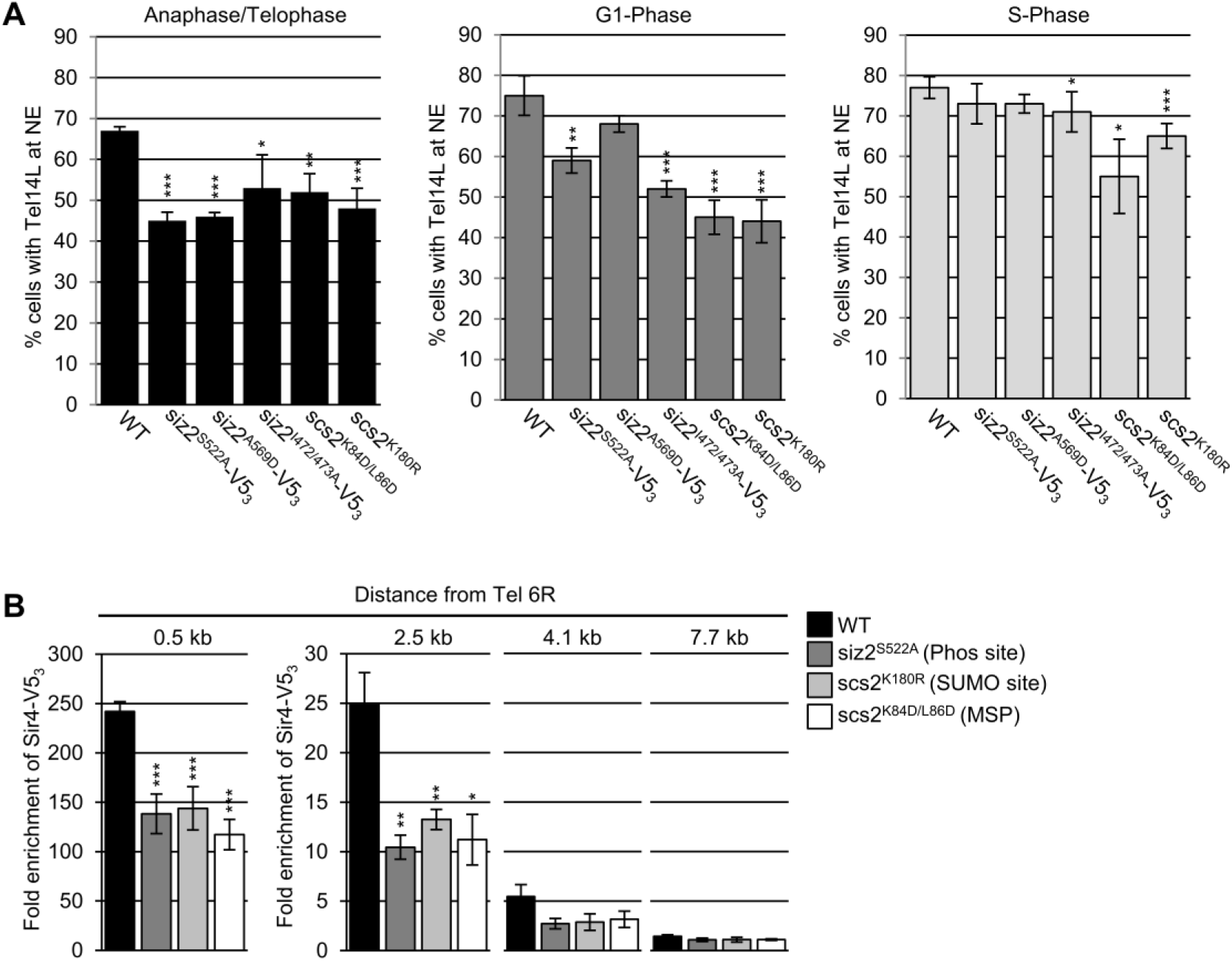
Telomere NE tethering during mitosis and G1-phase requires Scs2-Siz2 association and Scs2 SUMOylation. (**A**) Tethering of Tel14L to the NE was examined using epifluorescence imaging. The percentage of the total number of GFP-lacI/*Tel14L-LacO_256_* foci examined that overlapped with NE-associated Sec63-mCherry signal was determined for three or more biological replicates. The graph shows the average values for various strains at the indicated points in the cell cycle. Cell cycle stage was assessed by bud size and nuclear morphology. n = 50 cells/replicate/cell cycle stage. Error bars - SD. (**B**) Sir4-V5_3_ binding to chromatin adjacent to Tel6R was assessed by ChIP and qRT-PCR analysis using asynchronous cultures of the indicated strains. Graphs represent at least three biological replicates. Error bars - SEM. Asterisks - significant change relative to WT using a two-tailed student’s t-test. *p ≤ 0.05, **p ≤ 0.01, ***p ≤ 0.001.

Currently, the factors that contribute to telomere association with the INM during mitosis are unknown; however, various mechanisms have been described for telomere and subtelomeric chromatin tethering to the INM during other stages of the cell cycle. Key players include Sir4 and Yku70/80, which function to tether telomeres to the NE in G1- and S-phase (Taddei and Gasser, 2012; Kupiec, 2014). Sir4 and Yku80 are both SUMOylated by Siz2, with previous studies suggesting that Siz2-mediated SUMOylation directs Sir4, but not Yku80, dependent telomere tethering during G1(Ferreira et al., 2011). Similarly, we find that Sir4, but not Yku70/80, plays a significant role in M-phase telomere tethering (Fig. S3 C). Yet, cells lacking Siz2 showed no defects in the INM association or number of Sir4-GFP foci (Lapetina et al., 2017) (Fig. S3 D), suggesting Sir4 association with the INM is not altered in the absence of Siz2. We therefore tested the alternative possibility that the Scs2-Siz2 complex supports the association of Sir4 with subtelomeric chromatin by ChIP analyses of Sir4 at regions adjacent to Tel6R. Similar to previous studies (Van de Vosse et al., 2013; Moradi-Fard et al., 2016), analysis of WT cells revealed Sir4- V5_3_ bound near Tel6R, with the highest enrichment occurring near the telomere (0.5 kb), followed by a progressive decrease with increasing distance from the telomere (Fig. 5 B). In *siz2* and *scs2* mutant cells defective in mitotic SUMOylation, Sir4 enrichment was significantly reduced in all regions adjacent to Tel6R. By contrast, enhanced Scs2-Siz2 dependent mitotic SUMOylation (*ulp1^K352E/Y583H^-V5_3_*) did not alter levels of Sir4 bound near Tel6R (Fig. S3 E). Thus, the Scs2-Siz2 mitotic complex appears to mediate telomere tethering by supporting Sir4-subtelomeric chromatin association.

### SUMOylation stabilizes the association of Sir4 with subtelomeric chromatin in mitosis

Sir4 SUMOylation is dependent on Siz2 (Ferreira et al., 2011), suggesting that formation of the mitotic Scs2-Siz2 complex may be resoponsible for mediating Sir4-subtelomeric association through SUMOylation. As such, we assessed Sir4-SUMO levels by examining purified His_8_-SUMO conjugates isolated from strains producing Sir4-V5_3_. We found in *siz2^S522A^*, *scs2^K180R^*, or *scs2^K84D/L86D^* mutants, Sir4 SUMOylation was reduced as compared to WT cells (Fig. 6 A). By contrast, enhanced Scs2-Siz2 dependent mitotic SUMOylation (*ulp1^K352E/Y583H^-V5_3_*) showed increased SUMOylation of Sir4 (Fig. 6 B) and increased levels of SUMO at subtelomeric chromatin (Fig. S4 A). These observations indicate that the targeting of Siz2 to the INM during mitosis is required for proper Sir4 SUMOylation and the association of Sir4 with subtelomeric chromatin.

**Figure 6.**
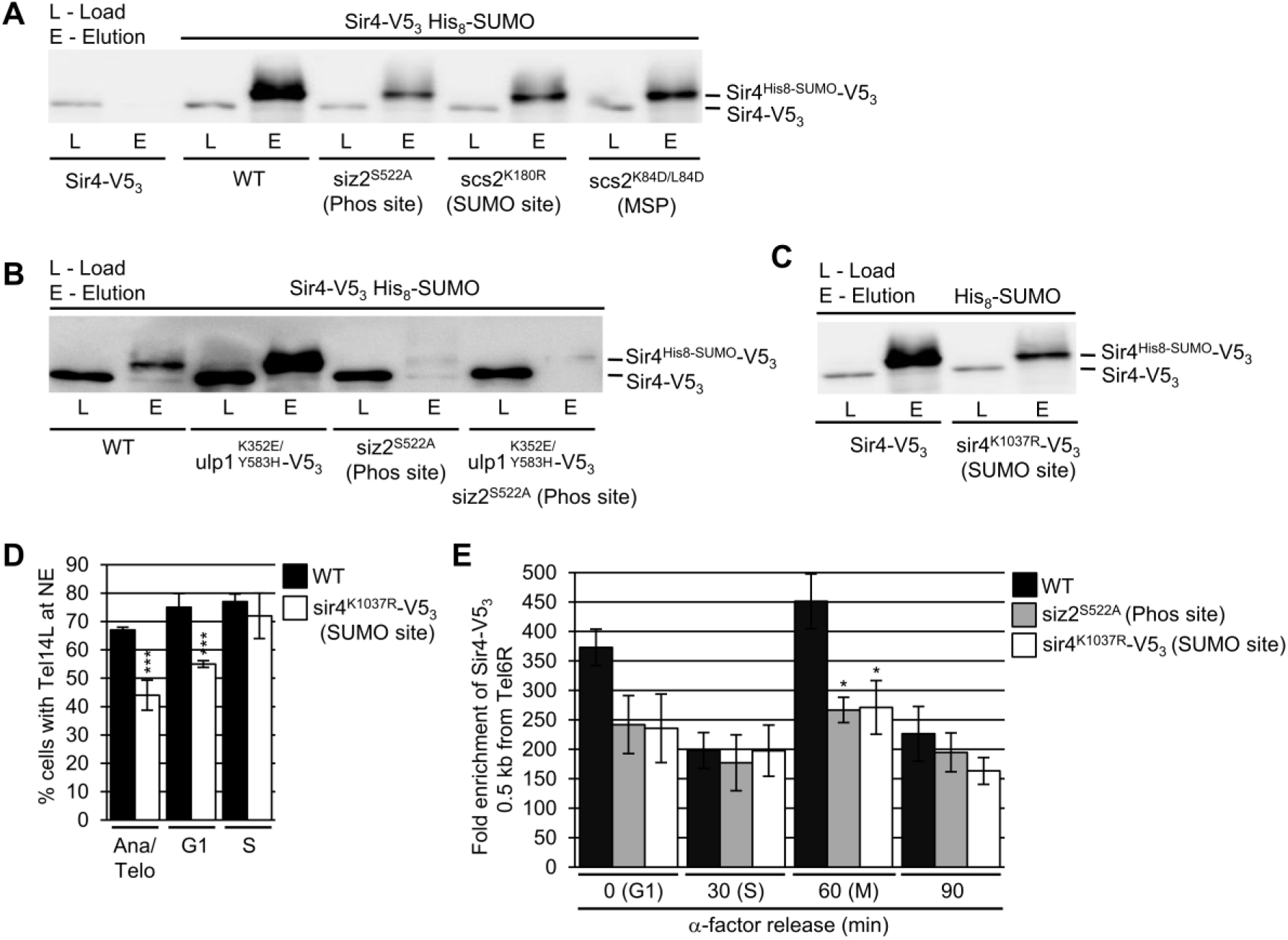
SUMOylation of Sir4 supports its incorporation into subtelomeric chromatin during mitosis. (**A-C**) His_8_-SUMO-conjugates were affinity-purified from the indicated strains containing Sir4-V5_3_ (**A**, **B**) or the sir4^K1037R^-V5_3_ SUMO site mutant (**C**). Levels of Sir4 and sir4^K1037R^ in the cell lysates (L), and Sir4-V5_3_-SUMO and sir4^K1037R^-V5_3_-SUMO in eluates (E) were examined by anti-V5 western blotting. (**D**) Tethering of Tel14L to the NE in the indicated strains was assessed as described in Fig. 5. WT data is the same as that used in Fig. 5 A. Graphs represent at least three biological replicates where n = 50 cells/replicate/cell cycle stage. Error bars - SD. (**E**) ChIP analysis was performed using synchronized cultures of the indicated strains at various cell cycle stages, including G1-phase (α-factor arrested cells), S-phase (30 min post α-factor release), and M-phase (60 min post α-factor release). Analysis was performed on a region 0.5 kb from Tel6R. Graphs represent at least three biological replicates. Error bars - SEM. Asterisks - significant change relative to WT using a two-tailed student’s t-test. *p ≤ 0.05, **p ≤ 0.01.

We further tested the significance of Sir4 SUMOylation by analyzing a Sir4 SUMOylation site mutant at residue K1037 within the Sir4 PAD domain (Zhao et al., 2014), a region necessary for telomere tethering activity (Andrulis et al., 2002; Taddei et al., 2004). The *sir4^K1037R^* mutant showed reduced Sir4 SUMOylation to levels similar to that detected in *siz2* and *scs2* point mutants (Fig. 6 A, 6 C). Moreover, NE tethering of telomeres Tel14L (Fig. 6 D) and Tel6R (Fig. S4 B) were also significantly reduced in the *sir4^K1037R^* mutant cells, specifically in M- and G1-phase nuclei, but not S-phase. These tethering defects did not appear to stem from mislocalization of Sir4 as NE association of the sir4^K1037R^-GFP mutant protein was indistinguishable from that of Sir4- GFP (Fig. S4 C). The sir4^K1037R^ mutant protein also showed a significantly reduced enrichment in all regions adjacent to Tel6R (Fig. S4 D). Notably, these phenotypes were similar to those observed in *siz2* and *scs2* point mutants exhibiting reduced Sir4 SUMOylation (Fig. 5 B, 6 A).

Finally, the defects in NE telomere tethering seen in both the *siz2^S522A^* and *sir4^K1037R^* mutant cells were restricted to M- and G1-phase (Fig. 5 A, 6 D and Fig. S4 D, S5 B). Therefore, we examined levels of Sir4-V5_3_ bound within a 0.5 kb region adjacent to Tel6R within synchronized cell cultures of these mutants (see Fig. S5) . In WT cells, this analysis revealed an ∼2-fold higher level of Sir4-V5_3_ associated with Tel6R in M- and G1-phase cells as compared to S-phase cells (Fig. 6 E). In cells where Sir4 SUMOylation is inhibited (*siz2^S522A^* or *sir4^K1037R^* mutant), S-phase levels of Sir4-bound Tel6R subtelomeric chromatin were similar to WT; however, in M- and G1- phase cells the association of Sir4 with Tel6R was reduced. These observations are consistent with a mitotic role for the Scs2-Siz2 complex in the SUMOylation of Sir4 and assembly of Sir4 into subtelomeric chromatin.

### NPC association of active *INO1* requires the Scs2-Siz2 complex

To extend our studies beyond telomeres, we examined the localization of the *INO1* gene under conditions of gene activation (e.g. growth in media lacking inositol). Previous studies have shown that *INO1* activation induces relocalization of the gene locus from the nucleoplasm to NPCs. This can be quantified by an increase in the frequency of cells displaying a NE-associated *INO1* locus (Brickner and Walter, 2004; Light et al., 2010). Our examination of the nuclear positioning of activated *INO1* during the cell cycle in WT cells revealed NE enrichment during M- and G1-phase (∼50-55% of cells), but not S-phase (Fig. 7), consistent with a previous report (Brickner and Brickner, 2010). However, in *siz2^S522A^* and *scs2^K180R^* point mutants, which inhibits SUMOylation of INM targets, the activated *INO1* locus did not show increased localization at the NE. In contrast, *INO1* localization appeared normal in *sir4^K1037R^* mutant cells. These observations indicate that the temporally and spatially regulated functions of the Scs2-Siz2 complex directs the establishment of different types of NE-chromatin interactions by multiple independent mechanisms.

**Figure 7.**
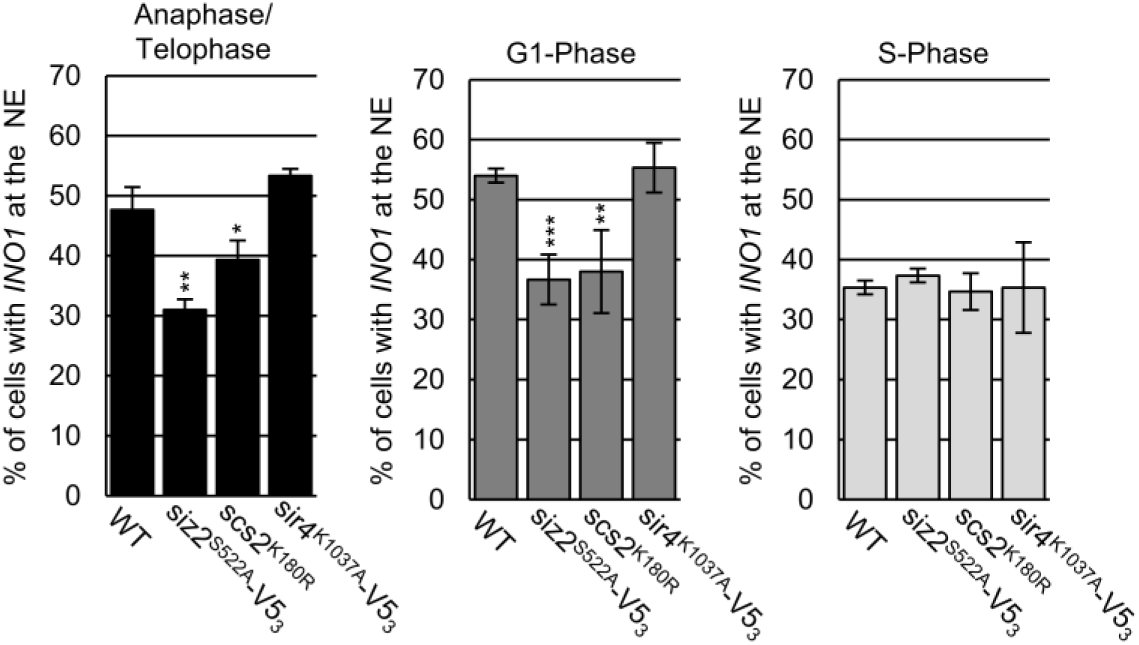
NPC association of activated *INO1* requires the Scs2-Siz2 complex and Scs2 SUMOylation. NE localization of the activated *INO1* locus was examined using epifluorescence imaging. The graphs show the percentage of total GFP-lac/*INO1-LacO_256_* foci that colocalize with NE localized Nup49-RFP. For each indicated strain, three biological replicates were assessed. Cell cycle stage was determined using bud size, and nuclear morphology. n = 50 cells/replicate/cell cycle stage. Error bars - SD. Asterisks - significant change relative to WT using a two-tailed student’s t-test. *p ≤ 0.05, **p ≤ 0.01, ***p ≤ 0.001.

## Discussion

As cells move from interphase into mitosis, chromatin disassociates from the INM in preparation for distribution of the genome to two postmitotic nuclei. Beginning in anaphase and into telophase, chromatin interactions with the NE membrane are reformed, establishing interactions of the INM and NPCs with chromatin (Ebrahimi and Donaldson, 2008; Brickner and Brickner, 2010; Poleshko et al., 2013; Poleshko et al., 2019). In this work, we show that a key step in this process in *S. cerevisiae* is the phosphorylation-dependent binding of the SUMO E3-ligase Siz2 to the INM-localized membrane protein Scs2. The Scs2-Siz2 complex initiates multiple mitotic SUMOylation events along the nucleoplasmic face of the NE that fosters reformation of NE-chromatin interactions and the faithful inheritance of epigenetic programs by newly forming nuclei.

### Spatiotemporal regulation of SUMOylation via Scs2

As cells progress through mitosis, we observed an accumulation of SUMO-conjugates along the NE that is directed by the phosphorylation of Siz2 and its subsequent NE association (Fig. 2). Our data suggest that Siz2 binding to the INM is, in part, controlled by a mechanism similar to that employed by Scs2 and other VAP protein family members to recruit cytoplasmic soluble and membrane-bound proteins to the ER (Brickner and Walter, 2004; Manford et al., 2012; Freyre et al., 2019). As in other cases (Goto et al., 2012; Kumagai et al., 2014; Weber-Boyvat et al., 2015; Kirmiz et al., 2018; Di Mattia et al., 2020), phosphorylation and an FFAT-like motif in Siz2 are critical for binding Scs2 (Fig. 2 and 3). However, unique to the Scs2-Siz2 complex is that its formation occurs in the nucleus and it uses a SUMO-SIM interaction to reinforce the FFAT-MSP domain interaction (Fig. 4). We envisage that a SUMO^Scs2^-SIM^Siz2^ interaction functions downstream of the MSP^Scs2^- FFAT/Phospho^-Siz2^ interaction to stabilize the Scs2-Siz2 complex at the INM.

As cells exit mitosis and progress through cytokinesis, the release of Siz2 from the INM is presumed to require dephosphorylation of Siz2 and deSUMOylation of Scs2. Our data led us to conclude that Scs2 deSUMOylation is performed by Ulp1 as strains expressing the *ulp1^K352E^* point mutation show dramatically increased levels of SUMOylated Scs2 during mitosis and into interphase of the next cell cycle (Fig. 1 D).

### A mitotic role for the Scs2-Siz2 complex in NE-chromatin interactions

The abundance of SUMO-conjugates along the INM during mitosis (Fig. 1 A) suggests numerous proteins and a broad spectrum of NE-associated processes are temporally regulated by these SUMOylation events. This high concentration of SUMO-modified proteins at the INM would establish a two-dimensional binding surface for SIM domain containing proteins. In this way, SUMO modifications at the INM could establish a multivalent interaction surface capable of promoting the assembly of various types of macromolecular complexes. Notably, the potential for SUMOylation to partition SIM-containing proteins near the INM parallels properties described in mammals for phase-separated PML bodies and their enhanced compartmentalization of SUMO and SIM-containing interacting partners (Van Damme et al., 2010; Banani et al, 2016; Min et al., 2019).

In this manner, as we show, the wave of mitotic NE SUMOylation supports interactions of both telomeres and the activated *INO1* gene with the NE during the later stages of mitosis (Fig. 5-7). With respect to telomeres, we further identified Sir4 as a key SUMOylation target of the Scs2-Siz2 complex (Fig. 6). The loss of Sir4 SUMOylation using various mutants (e.g., *siz2^S522A^* or *sir4^K1037R^*) causes reduced Sir4 binding to subtelomeric chromatin and lowers telomere binding to the INM during the later stages of mitosis and G1-phase of the subsequent cell cycle (Fig. 5, 6). For activated genes, we further note that the timing of Scs-Siz2 mediated SUMOylation correlates with the observed binding of several active genes, including *INO1*, with NPCs in M-phase following their dissociation from the NE in the preceding S-phase (Brickner and Brickner, 2010). As such, we envisage that Scs2-Siz2 directed SUMOylation broadly functions to support binding of multiple active gene loci to NPCs, including those employing transcription factors (Brickner et al., 2019), and especially those previously implicated to involve SUMOylation and Siz2 (Rosonina et al., 2010; Texari et al., 2013; Texari and Stutz, 2015; Saik et al., 2020).

Both the NE recruitment of Siz2 and SUMOylation of target proteins are transient, being largely reversed by G1-phase (Fig. 1, 2). We interpret this to suggest that SUMO-SIM interactions formed in mitosis guide and augment additional protein-protein interactions that are maintained in interphase and support chromatin-NE associations. Moreover, the removal of these SUMO-modifications may also play an important role. Interestingly, we observed that conditions that delay deSUMOylation of Scs2 and other INM proteins (*ulp1^K352E/Y583H^* mutant) led to increased INM retention of telomeres in G1-phase (Fig. S3 B). These results suggest that post-mitotic deSUMOylation may relax interactions and promote the periodic switching of telomeres between NE bound and unbound states observed in interphase cells (Hediger et al., 2002).

It is also important to consider that the multiple Scs2-Siz2 dependent SUMO-conjugates detected during mitosis may reflect a broader spectrum of NE-associated processes impacted by these temporal and spatial SUMOylation events. Scs2 in particular has roles in the regulation of lipid metabolism and the expression of genes controlling inositol biosynthesis (Loewen et al., 2003; Brickner and Walter, 2004; Manford et al., 2012; Gaspar et al., 2017). Consequently, it is possible that SUMOylation of Scs2 coupled with Siz2 recruitment could further regulate Scs2 functions, including lipid metabolism to accommodate NE expansion and the formation of two daughter nuclei. We expect future work aimed at the identification of INM targets of Scs2-Siz2, and the consequences of blocking these specific SUMOylation events, will be critical to addressing these possibilities.

## Methods

### Yeast strains and plasmids

Yeast strains are listed in Supplementary Table 1. Strains were derived from S288C (BY4741, BY4742) backgrounds except for telomere localization strains (W303) and *INO1* localization strains (YEF473A). Cells were grown in YPD (1% yeast extract, 2% bactopeptone, and 2% glucose) or synthetic medium (per L: 1.7g yeast nitrogen base, 5g ammonium acetate, 1.7 g amino acid dropout powder and 2% glucose), as required.

Transformations were performed using a lithium acetate/polyethylene glycol method (Gietz and Woods, 2002). Strains bearing gene modifications for protein fusions, gene deletions, and amino acid substitutions were generated using a PCR-based genomic integration method (Longtine et al., 1998), while strains bearing multiple gene modifications were derived by crossing relevant strains followed by sporulation and dissection. Deletion strains were generated by replacing the ORF of a given gene with a PCR cassette consisting of 60 bp 5’ of the ORF’s start codon - a marker gene - 60 bp 3’ of the ORF’s stop codon.

Coding sequence for the V5_3_ tag was integrated at the 3’end of *SIZ2*, *ULP1*, *SCS2*, and *SIR4* using the plasmid pTM1198 as previously described (Lapetina et al., 2017). Coding sequence for the HA_3_ tag was integrated at the 5’ end of *SCS2* using the plasmid pFA6a-kanMX6-SCS2pr- 3HA which was generated by replacing the *GAL1* promoter, bounded by BglII/PacI restriction enzyme sites in pFA6a-kanMX6-PGAL1-3HA, with 372 bp of the *SCS2* 5’UTR. Oligonucleotides used in PCRs to amplify *V5_3_* and *HA_3_* containing cassettes were designed as previously specified (Longtine et al., 1998).

Integration of the eGFP and PrA coding sequences at the 3’ end of *SIR4* were carried out as previously described (Lapetina et al., 2017). Scs2-TAP was obtained from the yeast TAP tag library (Ghaemmaghami et al., 2003). PCR cassettes used to integrate the coding sequence for mCherry at the 3’ end of *SUR4* and *NOP56*, were generated using the pGEM-4Z-mCherry-NAT plasmid as a template (Cairo et al., 2013).

pRS315.SMT3pr-His_8_-SMT3-HPH was constructed by cloning three PCR products into pRS315 (Sikorski and Hieter, 1989), including: 1) 350bp of the *SMT3* 5’ UTR bounded by SacI/NotI restriction enzyme sites; 2) the coding region for His_8_-Smt3 plus 324 bp of the *SMT3* 3’ UTR bounded by NotI/SalI restriction enzyme sites; 3) the *HPH-MX* sequence bounded by SalI/ApaI restriction sites. This plasmid was used as a PCR template, and the resulting SMT3pr- His_8_-SMT3-HPH cassette was integrated at the *SMT3* locus.

Amino acid substitutions were introduced into WT cells by site directed mutagenesis of genomic loci using a PCR-based one-step integration method. Genomic DNA derived from *SIR4-, SIZ2-, and ULP1-V5_3_* strains were used as the DNA templates. Sense oligonucleotides included mutagenic nucleotides within the relevant codon(s) of the target gene as well as ∼ 20 nucleotides downstream and ∼ 60 nucleotides upstream of this altered sequence to facilitate PCR amplification and genomic integration. Antisense oligonucleotides were the same as those used for C-terminal V5_3_ tagging. In the case of *scs2* mutants, genomic DNA from a *KAN-SCS2* strain, where the KAN marker was integrated between nucleotides 242/241 5’ of the *SCS2* start codon, was used as the PCR template. Antisense oligonucleotides included mutagenic nucleotides within the relevant *SCS2* codon(s), as well as ∼20 nucleotides downstream and ∼60 nucleotides upstream of this altered sequence to facilitate PCR amplification and genomic integration. The sense oligonucleotide encompassed 20 nucleotides within the 5’ end of the *KAN-MX TEF1* promoter and 60 bases upstream of position 242 5’ of the *SCS2* start codon. All mutations were sequence verified.

All GFP-siz2 fusions were generated in two steps. *GFP^S65T^* was integrated at the 5’ end of each *siz2-V5_3_-KAN* gene, as previously described for *SIZ2* (Lapetina et al., 2017). The *V5_3_-KAN* region was then replaced by integration of PCR cassette consisting of 60bp at the 3’ end of the *SIZ2* ORF - 273 bp after the *SIZ2* stop codon - *HPH-MX* - nucleotides 274-334 3’of the *SIZ2* stop codon. To generate *GFP-siz2^V720/721A^* the PCR cassette included the A720/721 codons. Regeneration of the 3’ end of the GFP-*siz2* genes was confirmed by sequencing. Similarly, the *V5_3_* coding sequence was removed from *siz2^S522A^-V5_3_-KAN* to generate *siz2^S522A^-NAT* strains. A similar strategy was used to remove the V5_3_ tag coding sequence and the Y583H mutation from *ulp1^K352E/Y583H^-V5_3_* to generate *ulp^K352E^*.

Strains used for split-superfolder GFP analysis were generated by first integrating a PCR cassette, consisting of *NAT-CDC42pr-GFP_1-10_* at the 5’ end of *SCS2, scs2^K180R^, and scs2^K84D/L86D^* using the previously described method (Smoyer et al., 2016). Plasmids encoding GFP_11_-mCherry- Hxk1 or GFP_11_-mCherry-Pus1 (Smoyer et al., 2016) where then transformed into the *NAT- CDC42pr-GFP_1-10_-SCS2, NAT-CDC42pr-GFP_1-10_- scs2^K180R^ NAT-CDC42pr-GFP_1-10_-scs2^K84D/L86D^* strains.

### α-factor arrest release assays

All strains used in α-factor arrest release assays were *MAT**a** bar1Δ*. 25 ml YPD cultures, incubated overnight at RT, were diluted to an OD_600_ = 0.3 in 30 ml of YPD followed by addition of α-factor to 10 ng/ml and incubation at 30°C for ∼ 2 hr 15 min to induce G1-phase arrest. Cells from an aliquot of this culture equivalent to OD_600_ = 1 were pelleted, resuspended in 50 μl of 2x SDS sample buffer and incubated at ∼ 80°C for 15 min. This represented the 0 min time point. The remaining cells were pelleted, washed once with 1 ml of YPD, pelleted again and resuspended in 25 ml fresh YPD to a final OD_600_ = 0.6, thereby removing α-factor. Cultures were grown at 30°C and sampled every ten minutes as described for the 0 min time point. All lysates were subsequently sonicated, and debris pelleted by centrifugation prior to running through an 8 % SDS-PAGE gel and subsequent western blot analysis. Blots were probed with the antibodies: anti-Smt3 (Wozniak Lab), anti-V5 (AbCam), anti-HA (Sigma), anti-Clb2 (Santa Cruz), and anti-Gsp1 (Wozniak Lab) as indicated.

For α-factor arrest release ChIP experiments, 50 ml cultures were incubated overnight at RT, then diluted to an OD_600_ of 0.2 in 500 ml of YPD and incubated for 1.5 hr at 30°C followed by addition of α-factor (10 ng/ml) and incubation at 30°C for ∼ 2 h 15 min. An OD_600_ = 50 equivalent was harvested from each culture, representing the 0 h time point, and treated as described for chromatin immunoprecipitation. To remove α-factor, the remaining cells were pelleted, washed twice with ddH_2_O and resuspended in 300 ml fresh YPD which was partitioned into three separate flasks. These cultures were incubated at 30°C for 30, 60 or 90 min prior to harvesting and subsequent treatment. Samples were then analyzed by chromatin immunoprecipitation as described below. OD_600_ = 1 equivalents were taken at all time points for western blot, as described above and for FACS analysis.

### Phosphatase treatment

To a 50 μl lysate sample, from the 60 min α-factor arrest-release time point using Siz2-V5_3_ cells, 200 μl of methanol, 50 μl of chloroform, and 150 μl of ddH_2_O were added sequentially, followed by vortexing for 10 s after each addition. The final mixture was centrifuged for 2 min at 15000 rpm. The resulting top layer was carefully removed, leaving the protein precipitate and the bottom layer. 300 μl of methanol was added to the remaining sample, and the mixture vortexed for 10 s followed by centrifugation for 2 min at 15000 rpm. Residual liquid was removed, and the resulting pellet air dried. The dried pellet was resuspended in 50 μl of 0.5% w/v SDS. To 10 ul of this sample 10 μl of lambda phosphatase buffer (NEB), 10 μl of 10 mM MnCl_2_, 1 μl lambda phosphatase (NEB) and 69 μl ddH_2_O was added. For the -PPase sample 1 μl lambda phosphatase was replaced with 1 μl ddH_2_O. Reactions were incubated at 30°C for 1 h, after which 20 µl of 50% TCA was added and the samples incubated overnight at 4°C. Samples were then centrifuged at 15 000 rpm for 20 min at 4°C, the supernatant removed and the remaining pellet air dried. The dried pellet was resuspended in 25 µl of 2x sample buffer and heated at 80°C for ∼15min prior to western blot analysis.

### *INO1* gene induction

Overnight synthetic media (+inositol) cultures, incubated at RT, were diluted into the same media to an OD_600_ = 0.2, then incubated at 30°C to an OD_600_ = ∼0.8. Cells were then pelleted, washed once with water, resuspended in synthetic media lacking inositol to an OD_600_ = 0.5 to induce *INO1* expression. Cultures were incubated at 30°C for 3 h followed by analysis by epifluorescence imaging as described below.

### Anti-SUMO immunofluorescnce

5 ml YPD cultures, incubated overnight at RT, were diluted to an OD_600_ = 0.2 in 5 ml of fresh YPD and incubated at 30°C to an OD_600_ ∼ 0.8. To each culture, 0.6 ml of 10X phosphate buffer (1M KH_2_PO_4_, 21g KOH per litre, 0.5 mM MgCl_2_) and 0.8 ml of 37% formaldehyde was added, followed by incubation at 30°C for 30 min. Cells were then pelleted and washed 2x with 1X phosphate buffer. The final pellets were resuspended in 100 μl of sorbitol-citrate buffer (100 mM K_2_PO_4_, 3.6 mM citric acid, 1.2 M sorbitol, 0.5 mM MgCl_2_) and DTT added to final concentration of 1 mM. Cells were then pelleted and resuspended in 100 μl of sorbitol-citrate buffer supplemented with 2 mg/ml 20T zymolyase and incubated at 30°C for 20 min. Cells were then pelleted, washed 2X with 1 ml sorbitol-citrate buffer, and the final pellet resuspended in 50 μl of sorbitol-citrate buffer. 20 μl of the cell suspension was pipetted onto a multi-well slide coated with 0.1 % poly-L-lysine and the slide incubated at RT in a covered box lined with damp paper towels for 30 min. All subsequent incubations and washes were carried out at RT in the same box. ∼ 20 μl of solution was used at each step, and solutions were exchanged by their aspiration off of the wells followed by the addition of fresh solution. Steps included 1)1x PBST wash, 2) 1x addition of PBS 0.1% Triton X-100 with a 10 min incubation, 3) 2x wash with PBST 4) 1x addition of PBST, 1% BSA with a 10 min incubation, 4) 1x addition of PBST, 1% BSA supplemented with an anti-SUMO antibody at a 1:200 dilution with a 1 h incubation, 5) 10x wash with PBST, 0.1% BSA 6) 1x addition of PBST, 1% BSA supplemented with Alexa Flouor 488 donkey anti-rabbit IgG antibody (Life technologies) at a 1:200 dilution, 6) 10x wash with PBST, 0.1% BSA. After the final wash, ∼ 3 ul of DAPI-Fluoromount-G (SouthernBiotech) was added to each well and a coverslip placed over the slide. Cells were then analyzed by epifluorescence imaging.

### Epifluorescence imaging and analysis

For live cell imaging of GFP-Siz2, split superfolder-GFP, telomere tagged, Sir4-GFP strains, YPD or synthetic media cultures, incubated overnight at RT, were diluted to an OD_600_ ∼ 0.1, then incubated at 30°C to an OD_600_ ∼ 1, except for GFP-Siz2 strains which were incubated at RT. Telomere tagged strains were also grown in YPD supplemented with 120 μg/ml of adenine. Cells from 1 ml of culture was then pelleted, washed with 1 ml of synthetic complete medium, and the result pellet suspended in ∼ 20 μl of synthetic complete medium. 1.5 μl of this cell suspension was then spotted onto a microscope slide coated with a 1% agarose pad consisting of 1g of agarose per 100 ml synthetic complete medium.

Epifluorescence images of cells prepared for anti-SUMO immunofluorescence, as well as cells producing GFP-Siz2 derivatives or split-superfolder-GFP fusions, were acquired using an Axio Observer.Z1 microscope (Carl Zeiss Inc.), equipped with an UPlanS-Apochromat 100x/1.40 NA oil objective lens (Carl Zeiss Inc.) and an AxioCam MRm digital camera with a charge-coupled device (Carl Zeiss Inc.). Images were saved using the AxioVision software and rendered using Image J software (National Institute of Health) for display. The unsharp mask filter was used to analyze anti-SUMO immunofluorescence images (Radius (Sigma): 2.0 pixels; Mask Weight: 0.7) as well as GFP-Siz2 and Sur4-mCherry images (Radius (Sigma): 3.0 pixels; Mask Weight: 0.8).

Epifluorescence images of cells containing GFP tagged telomeres, Sir4-eGFP, or GFP tagged *INO1* were acquired on a DeltaVision Elite imaging system (GE Healthcare Life Sciences) with a 60x/1.42 NA oil, Plan Apo N objective (Olympus). Images were collected as 15 x 0.2 μm z-stacks using the SoftWoRx software, (version 6.5.2, GE Healthcare Life Sciences), then rendered and analyzed using Image J (NIH).

Cell cycle stage was assessed based on bud size and/or nuclear morphology. Specifically, G1 phase -unbudded cells, and S phase-small-budded cells, anaphase/telophase - barbell shaped nuclei. Localization of foci corresponding to Sir4-eGFP, GFP tagged *INO1*, and GFP tagged telomeres in anaphase/telophase cells, as well foci corresponding to GFP tagged telomeres in G1- and S-phase of in *ulp1^K352E/Y583H^-V5_3_* cells and corresponding WT cells (Fig. S4), were scored positive when the focus fully or partially overlapped with the NE localized marker (i.e. Sur4- mCherry, Nup49-RFP, or Sec63-GFP). Subnuclear localization of G1- and S-phase foci of GFP-tagged telomeres was determined, as previosuly described (Lapetina et al., 2017), by dividing the telomere distance from the NE (TD) by the nuclear radius (*r*). The TD/*r* ratio (R) was used to group telomeres into three concentric zones of equal area. Zone 1 represents foci with ratios ≤ 0.184 x R; zone 2 foci with ratios > 0.184 x R and < 0.422 x R; and zone 3 represents foci with ratios ≥ 0.422 x R. This method was only used in cells, and at cell cycle stages where the nuclei remained spherical.

For all cell cycle dependent imaging experiments the number of foci associated with the NE was expressed as a percentage of the total number of foci observed. At least three replicates were assessed for each strain, where n = 50 cells/replicate/cell cycle stage. Error bars in these experiments represent SD, and significance was assessed relative to a WT control using a two-tailed student’s t-test where *p ≤ 0.05, **p ≤ 0.01, ***p ≤ 0.001.

### Western blot analysis

Proteins separated using SDS-PAGE were transferred to nitrocellulose membranes. Membranes were incubated in blocking buffer (TBST or PBST with 5% milk powder) for at least 1 h at RT. Fresh blocking buffer supplemented with primary antibody was then added, followed by incubation overnight at 4°C. Membranes were then washed 3X with TBST or PBST, then incubated in fresh blocking buffer supplemented with a secondary antibody-HRP conjugate (BioRad) at a 1:10000 dilution for at least 1 h at RT. Membranes were washed 3X with TBST or PBST and proteins were visualized by chemiluminescence using an ImageQuant LAS 4000 (GE) imaging system. All western blot images were rendered using Image J software (National Institute of Health).

### Coaffinity Purification

Coaffinity purifications were carried as previously described (Van de Vosse et al., 2013). 50 ml YPD cultures of Scs2-TAP producing cells, grown overnight at RT, were diluted in 1 l of fresh YPD to an OD_600_ ∼ 0.1, then incubated at 30°C to an OD_600_ ∼ 1.0. Cells were pelleted, washed once with 25 ml cold ddH_2_O, and the resulting pellet extruded through a 5 ml syringe directly into a 50 ml falcon tube containing liquid nitrogen producing cell noodles. Liquid nitrogen was removed, and the noodles stored at -80°C. Noodles were then subjected to at least seven rounds of ball mill grinding (Reitch PM100; 1 min 15s, 450 rpm per round) keeping the grinding vessel cold between rounds by partial immersion in liquid nitrogen. The resulting cell powder was stored at -80°C.

To 1 g of cell powder, 2ml of IP buffer (2mM MgCl_2_, 20mM HEPES-KOH (pH 7.4), 0.1% Tween-20, 110mM KOAc, antifoam-B emulsion at 1:5000 dilution, and protease inhibitors (2 complete EDTA-free pellets (Roche)/50 ml buffer) was added. The suspension was incubated on ice for 30 min, with vortexing every 5 min. The resulting lysate was cleared by centrifugation at 1 500 g for 10 min at 4°C. 25 µl of the clarified lysate, representing the load, was added to 1 ml of ddH_2_0, followed by TCA precipitation, and resuspension of the resulting pellet in 75 μl of 2X sample buffer. 3 mg of IgG-conjugated magnetic beads (Dynabeads; Invitrogen) in 100 μl of IP buffer was added to the 2 ml of clarified cell lysate, and the mixture was incubated for 1 h at 4°C with rotation. Beads were collected using a magnet and washed 10x with 1ml of IP buffer at 4°C. 1ml of the last wash was collected and subjected to TCA precipitation as described. Proteins bound to beads were eluted at 4°C using 0.5 ml IP buffer containing incrementally increasing concentrations of MgCl_2_ (0.05, 0.5, and 2 M) followed by a final elution using 0.5 ml of 0.5 M acetic acid to release the TAP fusion protein from the beads. To the 500 µl eluate fractions 500 µl of ddH_2_0 was added followed by TCA precipitation as described. All samples collected were analyzed by western blotting.

### His_8_-SUMO pulldowns

Frozen cell powders derived from strains producing Sir4-V5_3_ and His_8_- Smt3 (His_8_-SUMO) were produced as described for coaffinity purification. Note that all 8M urea/phosphate buffers were made fresh just prior to use. To 1 g of frozen cell powder, 10 ml of resuspension buffer (8 M Urea, 50 mM NaPO_4_ (pH 8.0), 500 mM NaCl, 1% NP40-Igepal) was added and the mixture incubated at RT with intermittent vortexing. Lysates were centrifuged at 15 000 rpm for 20 min. 25 μl of the clarified lysate, was added to 1 ml ddH_2_0, followed by TCA precipitation and resuspension in 100 μl of 2X sample buffer (Load sample). The remaining lysate was transferred to a 15 ml falcon tube and 1 ml of a 50% slurry consisting of NiNTA agarose beads (Qiagen) in resuspension buffer was added per 10 ml of lysate. The mixture was incubated at RT with rotation for 2 h. NiNTA agarose beads were then pelleted by centrifugation at 1 000 rpm for 1 min and the lysate removed. The beads were then washed 3X with 6X bead volume of wash buffer (8 M Urea, 50 mM NaPO_4_ (pH 6.3), 500 mM NaCl, 1% NP40-Igepal). After the last wash, 2 ml of elution buffer (8 M Urea, 50 mM NaPO_4_, 500 mM NaCl, 1% NP40-Igepal adjusted to pH 4.5) was added to the beads and the slurry incubated at RT with rotation for 1 h followed by centrifugation at 1 000 rpm for 1 min. The resulting eluant was collected and subjected to TCA precipitation, with samples resuspended in a final of 100 μl of 2X sample buffer. Load and eluate samples were then analyzed by western blotting.

### Chromatin Immunoprecipitation

10 ml YPD cultures, incubated overnight at RT, were diluted into 50 ml of fresh YPD to an OD_600_ ∼ 0.2 and grown at 30°C to an OD_600_ ∼ 0.8. Cells from an OD_600_ = 50 equivalent of each culture was pelleted, and the cells resuspended in 50 ml of YPD 1% formaldehyde, followed by incubation at 30°C for 20 min to induce crosslinking. Glycine was added to 125mM followed by a 5 min incubation at RT to quench crosslinking. Cells were pelleted, washed with TBS and the resulting pellet flash frozen using liquid nitrogen and stored at -80°C.

Cells were resuspended in 500 μl FA lysis buffer (50mM HEPES-KOH pH 7.5, 150mM NaCl, 1mM EDTA, 1% Triton X-100, 0.1% Na deoxycholate) and disrupted by glass bead lysis at 4°C. Glass beads were removed from the lysate, and the lysates then subjected to sonication, while keeping samples on ice, to shear chromosomal DNA to an average size of ∼ 400bp, as assessed by electrophoresis. Lysates were diluted by the addition of FA lysis buffer to a final volume of 1.5 ml. 500 μl, representing the Input, was then diluted 100X into a TE 1% SDS solution, and subsequently reverse crosslinked as described below. The remaining lysate was incubated with 5 μl of ssDNA (10mg/ml) and 2 µl of mouse monoclonal anti-V5 antibody (Abcam), 4 µl of rabbit polyclonal anti-PrA (Sigma) antibody or 4 µl of rabbit polyclonal anti-Smt3 (SUMO) antibody (Wozniak lab) for 2 hours at 4°C. 30 µl of Protein G Dynabeads (Invitrogen) in FA lysis buffer was added to each lysate followed by 1 h incubation at 4°C. Beads were collected by magnet and sequentially washed 2x with FA lysis buffer, 1x with FA lysis buffer including 500mM NaCl, 1x with wash buffer (10mM Tris-HCl pH 8, 0.25M LiCl, 1mM EDTA, 0.5% NP-40, 0.5% sodium deoxycholate), and 1X with TE buffer. After the final wash, chromatin was eluted from the beads using 2 rounds of incubation at 65°C for 10 minutes with 200 μl of TE 1% SDS. Input and ChIP samples were then reverse cross-linked by overnight incubation at 65°C, followed by addition of 5 µl of Proteinase K (20mg/ml) and 1 µl of glycogen (20mg/ml) to the samples and incubation at 37°C for 2 h. 40 µl of 5M LiCl was then added to each sample, followed by phenol/chloroform extraction and ethanol precipitation. The resulting DNA pellets were resuspended in 50 µl TE and 5 µl of RNase A (5mg/ml) then incubated at 37°C for 1 h followed by purification using Qiagen PCR Purification Kit.

Samples were analyzed by RT-qPCR as previously described (Makio and Wozniak, 2020). ChIP and Input DNA were used to amplify target sequences of interest using PerfeCTa SYBR green PCR mix (Quanta Biosciences) and a MX3000 (Agilent) instrument. The relative fold enrichment of the subtelomeric chromatin immunoprecipitated with the protein of interest was evaluated by the ΔΔCt method (Livak and Schmittgen, 2001). PCR amplification of each subtelomeric region was first normalized against the amplification of the corresponding input DNA to generate a ΔCt value. Each subtelomeric region was then normalized to the amplification of a non-specific binding control region including either 17.1 kb from Tel6R (Sir4 ChIP) or an intergenic region in Chromosome V (SUMO ChIP), to generate ΔΔCt values. The relative fold enrichment of subtelomeric chromatin over background was given as 2^-ΔΔCt^, based on the assumption that the PCR reaction was 100% efficient. Sense (S) and antisense (AS) primers used for qRT-PCR included: Chromosome V intergenic region (S-ACATTCTTGGAAACCCATCG), (AS-TCGTATCATGAT TTAGCGTCGT); Chr. VIR: 0.5 kb (S- GATAACTCTGAACTGTGCATCCAC), (AS-ACTGTC GGAGAGTTAACAAGCGGC); 2.5 kb (S-GAGCAATGAATCTTCGGTGCTTGG), (AS-CGCAGTACCTTGGAAAAATCTAGGC); 4.1 kb (S-CGTTCTTCTTGGCCCTTATC), (AS-CATCATCGGTGGTTTTGTCGTG); 7.7 kb (S- AAGTCACTATGGGTTGCCGGTATC), (AS-AACT ACCTCTATAGGACCTGTCTC); 17.1 kb (S-GAAAGTTTGGATGCTAGCAAG GGC), (AS-GCATAGCCTTTGAAAACGGCG).

### FACS

Cells pelleted for DNA content determination by FACS (see α-factor arrest release assays) were resuspended in 1 ml of 70% ethanol and incubated overnight at 4°C. Cells were then pelleted and the pellets washed 2X with 1 ml 50 mM Tris-HCl (pH 8.0). Final pellets were resuspended in 0.5 ml of 50 mM Tris-HCl (pH 8.0) containing RNase A (0.4 mg/ml) and incubated at 37°C for 2 h. Cells were then washed 2X with 1 ml 50 mM Tris-HCl (pH 8.0) and the final pellets resuspended in 200 μl of a 5 mg/ml pepsin solution (5 mg pepsin, 5 ul conc. HCl, per ml ddH_2_0), and incubated at 37°C for 1 h. 1 ml of 50 mM Tris-HCl (pH 8.0) was added, and the cells were pelleted then washed once with 1 ml of the same buffer. Post wash, pellets were suspended in 250 μl propidium iodide solution (50 mM Tris-HCl (pH 8.0), 50 μg/ml propidium iodide) and incubated overnight at 4°C. 50 μl of this cell suspension was added to 2 ml of 50 mM Tris-HCl (pH 8.0) in a round bottom tube and the sample briefly sonicated at low power to resuspend cells. DNA content was then determined using a BD LSRFortessa cell analyzer (Software Version 2.0).

## Supporting information

All supplementary material

## Acknowledgments

We thank Dr. Michael Hendzel (University of Alberta), and members of the Aitchison, Montpetit, and Wozniak labs for helpful discussions. Funding for this work is supported by the Canadian Institutes of Health Research (MOP 106502 and 36519) to RWW and the National Institutes of Health, USA (NCDIR: 2P41GM109824-06 and R01: 2R01GM112108-05 to JDA and R01GM124120 to BM). The authors declare no competing financial interests.

## Notes

### Competing Interest Statement

The authors have declared no competing interest.

## References

Albuquerque, C.P., M.B. Smolka, S.H. Payne, V. Bafna, J. Eng, and H.A. Zhou. 2008. A multidimensional chromatography technology for in-depth phosphoproteome analysis. Mol. Cell. Proteom. 7:1389–1396. doi:10.1074/mcp.M700468-MCP200.

Andrulis, E.D., D.C. Zappulla, A. Ansari, S. Perrod, C.V. Laiosa, M.R. Gartenberg, and R. Sternglanz. 2002. Esc1, a nuclear periphery protein required for Sir4-based plasmid anchoring and partitioning. Mol Cell Biol. 22:8292–8301. doi:10.1128/mcb.22.23.8292-8301.2002.

Banani, S.F., A.M., Rice, W.B. Peeples, Y. Lin, S. Jain, R. Parker, and M.K. Rosen. 2016. Compositional control of phase-separated cellular bodies. Cell. 166:651–663. doi:10.1016/j.cell.2016.06.010.

Brickner, J.H., and P. Walter. 2004. Gene recruitment of the activated INO1 locus to the nuclear membrane. PLoS Biol. 2:e342. doi:10.1371/journal.pbio.0020342.

Brickner, D.G., and J.H. Brickner. 2010. Cdk phosphorylation of a nucleoporin controls localization of active genes through the cell cycle. Mol. Biol. Cell. 21:3421–3432. doi:10.1091/mbc.E10-01-0065.

Brickner, D.G., C. Randise-Hinchliff, M. Lebrun Corbin, J.M. Liang, S. Kim, B. Sump, A. D’Urso, S.H. Kim, A. Satomura, H. Schmit, R. Coukos, S. Hwang, R. Watson, and J.H. Brickner. 2019. The role of transcription factors and nuclear pore proteins in controlling the spatial organization of the yeast genome. Dev. Cell. 49:936–947. doi:10.1016/j.devcel.2019.05.023.

Buchwalter, A., J.M. Kaneshiro, and M.W. Hetzer. 2019. Coaching from the sidelines: the nuclear periphery in genome regulation. Nat. Rev. Genet. 20:39–50. doi:10.1038/s41576-018-0063-5.

Cairo, L.V., C. Ptak and R.W. Wozniak. 2013. Mitosis-specific regulation of nuclear transport by the spindle assembly checkpoint protein Mad1p. Mol. Cell. 49:109–120. doi:10.1016/j.molcel.2012.10.017.

Champion, L., S. Pawar, N. Luithle, R. Ungricht, and U. Kutay. 2019. Dissociation of membrane-chromatin contacts is required for proper chromosome segregation in mitosis. Mol. Biol. Cell. 30:427–440. doi:10.1091/mbc.E18-10-0609.

Chao, J.T., A.K. Wong, S. Tavassoli, B.P. Young, A. Chruscicki, N.N. Fang, L.J. Howe, T. Mayor, L.J. Foster, and C.J. Loewen. 2014. Polarization of the endoplasmic reticulum by ER-septin tethering. Cell. 158:620–632. doi:10.1016/j.cell.2014.06.033.

Cuijpers, S.A.G, and A.C.O. Vertegaal. 2018. Guiding mitotic progression by crosstalk between post-translational modifications. Trends Biochem. Sci. 43:251–268. doi:10.1016/j.tibs.2018.02.004.

Denison C., A.D. Rudner, S.A. Gerber, C.E. Bakalarski, D. Moazed, and S.P. Gygi. 2005. A proteomic strategy for gaining insights into protein sumoylation in yeast. Mol. Cell Proteomics. 4:246–254. doi:10.1074/mcp.M400154-MCP200.

Di Mattia, T., A. Martinet, S. Ikhlef, A.G. McEwen, Y. Nominé, C. Wendling, P. Poussin-Courmontagne, L. Voilquin, P. Eberling, F. Ruffenach, J. Cavarelli, J. Slee, T.P. Levine, G. Drin, C. Tomasetto, and F. Alpy. 2020. FFAT motif phosphorylation controls formation and lipid transfer function of inter-organelle contacts. EMBO J. 39:e104369. doi:10.15252/embj.2019104369.

Ebrahimi, H., and A.D. Donaldson. 2008. Release of yeast telomeres from the nuclear periphery is triggered by replication and maintained by suppression of Ku-mediated anchoring. Genes Dev. 22:3363–3374. doi:10.1101/gad.486208.

Egecioglu, D., and J.H. Brickner. 2011. Gene positioning and expression. Curr. Opin. Cell Biol. 23:338–345. doi:10.1016/j.ceb.2011.01.001.

Encinar Del Dedo, J., F.Z. Idrissi, I.M. Fernandez-Golbano, P. Garcia, E. Rebollo, M.K. Krzyzanowski, H. Grotsch, and M.I. Geli. 2017. ORP-mediated ER contact with endocytic sites facilitates actin polymerization. Dev. Cell. 43:588–602. doi:10.1016/j.devcel.2017.10.031.

Falk, M., Y. Feodorova, N. Naumova, M. Imakaev, B.R. Lajoie, H. Leonhardt, B. Joffe, J. Dekker, G. Fudenberg, I. Solovei, and L.A. Mirny. 2019. Heterochromatin drives compartmentalization of inverted and conventional nuclei. Nature. 570:395–399. doi:10.1038/s41586-019-1275-3.

Felberbaum, R., N.R. Wilson, D. Cheng, J. Peng, and M. Hochstrasser. 2012. Desumoylation of the endoplasmic reticulum membrane VAP family protein Scs2 by Ulp1 and sumo regulation of the inositol synthesis pathway. Mol. Cell. Biol. 32:64–75. doi:10.1128/MCB.05878-11.

Ferreira, H.C., B. Luke, H. Schober, V. Kalck, J. Lingner, and S.M. Gasser. 2011. The PIAS homologue Siz2 regulates perinuclear telomere position and telomerase activity in budding yeast. Nat. Cell Biol. 13:867–874. doi:10.1038/ncb2263.

Freudenreich, C.H., and X.A. Su. 2016. Relocalization of DNA lesions to the nuclear pore complex. FEMS Yeast Res. 16:fow095. doi:10.1093/femsyr/fow095.

Freyre, C.A.C., P.C. Rauher, C.S. Ejsing, and R.W. Klemm. 2019. MIGA2 links mitochondria, the ER, and lipid droplets and promotes de novo lipogenesis in adipocytes. Mol. Cell. 76:811–825. doi:10.1016/j.molcel.2019.09.011.

Gaspar, M.L., Y.F. Chang, S.A. Jesch, M. Aregullin, and S.A. Henry. 2017. Interaction between repressor Opi1p and ER membrane protein Scs2p facilitates transit of phosphatidic acid from the ER to mitochondria and is essential for INO1 gene expression in the presence of choline. J. Biol. Chem. 292:18713–18728. doi:10.1074/jbc.M117.809970.

Gietz, R.D., and R.A. Woods. 2002. Transformation of yeast by lithium acetate/single-stranded carrier DNA/polyethylene glycol method. Methods Enzymol. 350:87–96. doi:10.1016/s0076-6879(02)50957-5.

Ghaemmaghami, S., W.K. Huh, K. Bower, R.W. Howson, A. Belle, N. Dephoure, E.K. O’Shea, and J.S. Weissman, J.S. 2003. Global analysis of protein expression in yeast. Nature. 425:737– 741. doi:10.1038/nature02046.

Goto, A., X. Liu, C.A. Robinson, and N.D. Ridgway. 2012. Multisite phosphorylation of oxysterol-binding protein regulates sterol binding and activation of sphingomyelin synthesis. Mol. Biol. Cell. 23:3624–3635. doi:10.1091/mbc.E12-04-0283.

Güttinger, S., E. Laurell, and U. Kutay. 2009. Orchestrating nuclear envelope disassembly and reassembly during mitosis. Nat. Rev. Mol. Cell Biol. 10:178–191. doi:10.1038/nrm2641. PMID: 19234477.

Hannich, J.T., A. Lewis, M.B. Kroetz, S.J. Li, H. Heide, A. Emili, and M. Hochstrasser. 2005. Defining the sumo-modified proteome by multiple approaches in Saccharomyces cerevisiae. J. Biol. Chem. 280:4102–4110. doi:10.1074/jbc.M413209200.

Hari, K.L., K.R. Cook, and G.H. Karpen. 2001. The Drosophila Su(var)2-10 locus regulates chromosome structure and function and encodes a member of the PIAS protein family. Genes Dev. 15:1334–1348. doi:10.1101/gad.877901.

Hediger, F., F.R. Neumann, G. Van Houwe, K. Dubrana, and S.M. Gasser. 2002. Live imaging of telomeres: yKu and Sir proteins define redundant telomere-anchoring pathways in yeast. Curr. Biol. 12:2076–2089. doi:10.1016/s0960-9822(02)01338-6.

Holt, L.J., B.B. Tuch, J. Villen, A.D. Johnson, S.P. Gygi, and D.O. Morgan. 2009. Global analysis of Cdk1 substrate phosphorylation sites provides insights into evolution. Science. 325:1682–1686. doi:10.1126/science.1172867.

Irniger, S. 2002. Cyclin destruction in mitosis: a crucial task of Cdc20. FEBS Lett. 532:7–11. doi:10.1016/s0014-5793(02)03657-8.

Jentsch, S., and I. Psakhye. 2013. Control of nuclear activities by substrate-selective and protein-group sumoylation. Annu. Rev. Genet. 47:167–186. doi:10.1146/annurev-genet-111212-133453.

Johnson, E.S., and A.A. Gupta. 2001. An E3-like factor that promotes sumo conjugation to the yeast septins. Cell. 106:735–744. doi:10.1016/s0092-8674(01)00491-3.

Kaiser, S.E., J.H. Brickner, A.R. Reilein, T.D. Fenn, P. Walter, and A.T. Brunger. 2005. Structural basis of FFAT motif-mediated ER targeting. Structure. 13:1035–1045. doi:10.1016/j.str.2005.04.010.

Kirmiz, M., N.C. Vierra, S. Palacio, and J.S. Trimmer. 2018. Identification of VAPA and VAPB as Kv2 channel-interacting proteins defining endoplasmic reticulum-plasma membrane junctions in mammalian brain neurons. J. Neurosci. 38:7562–7584. doi:10.1523/JNEUROSCI.0893-18.2018.

Kumagai, K., M. Kawano-Kawada, and K. Hanada. 2014. Phosphoregulation of the ceramide transport protein CERT at serine 315 in the interaction with VAMP-associated protein (VAP) for inter-organelle trafficking of ceramide in mammalian cells. J. Biol. Chem. 289:10748–10760. doi:10.1074/jbc.M113.528380.

Kupiec, M. 2014. Biology of telomeres: lessons from budding yeast. FEMS Microbiol. Rev. 38:144–171. doi:10.1111/1574-6976.12054.

Lapetina, D.L., C. Ptak, U.K. Roesner, and R.W. Wozniak, R.W. 2017. Yeast silencing factor Sir4 and a subset of nucleoporins form a complex distinct from nuclear pore complexes. J. Cell. Biol. 216:3145–3159. doi:10.1083/jcb.201609049.

Light, W.H., D.G. Brickner, V.R. Brand, and J.H. Brickner. 2010. Interaction of a DNA zip code with the nuclear pore complex promotes H2A.Z incorporation and INO1 transcriptional memory. Mol. Cell. 40:112–125. doi:10.1016/j.molcel.2010.09.007.

Livak, K.J., and T.D. Schmittgen. 2001. Analysis of relative gene expression data using real-time quantitative PCR and the 2(-Delta Delta C(T)) method. Methods. 25:402–408. doi:10.1006/meth.2001.1262.

Loewen, C.J., A. Roy, and T.P. Levine. 2003. A conserved ER targeting motif in three families of lipid binding proteins and in Opi1p binds VAP. EMBO J. 22:2025–2035. doi:10.1093/emboj/cdg201.

Loewen, C.J., and T.P. Levine. 2005. A highly conserved binding site in vesicle-associated membrane protein-associated protein (VAP) for the FFAT motif of lipid-binding proteins. J. Biol. Chem. 280:14097–14104. doi:10.1074/jbc.M500147200.

Loewen, C.J., B.P. Young, S. Tavassoli, and T.P. Levine. 2007. Inheritance of cortical ER in yeast is required for normal septin organization. J. Cell Biol. 179:467–483. doi:10.1083/jcb.200708205.

Longtine, M.S., A. McKenzie, D.J. Demarini, N.G. Shah, A. Wach, A. Brachat, P. Philippsen, and J.R. Pringle. 1998. Additional modules for versatile and economical PCR-based gene deletion and modification in Saccharomyces cerevisiae. Yeast. 14:953–961. doi:10.1002/(SICI)1097-0061(199807)14:10<953::AID-YEA293>3.0.CO;2-U. PMID: 9717241.

Ma, Y., K. Kanakousaki, and L. Buttitta. 2015. How the cell cycle impacts chromatin architecture and influences cell fate. Front. Genet. 6:19. doi:10.3389/fgene.2015.00019.

Manford, A.G., C.J. Stefan, H.L. Yuan, J.A. Macgurn, and S.D. Emr. 2012. ER-to-plasma membrane tethering proteins regulate cell signaling and ER morphology. Dev. Cell. 23:1129–1140. doi:10.1016/j.devcel.2012.11.004.

Makio, T., and R.W. Wozniak. 2020. Passive diffusion through nuclear pore complexes regulates levels of the yeast SAGA and SLIK coactivator complexes. J. Cell Sci. 133:jcs237156. doi:10.1242/jcs.237156.

Min, J., W.E. Wright, and J.W. Shay. 2019. Clustered telomeres in phase-separated nuclear condensates engage mitotic DNA synthesis through BLM and RAD52. Genes Dev. 33:814–827. doi:10.1101/gad.324905.119.

Misteli, T. 2020. The Self-Organizing Genome: Principles of Genome Architecture and Function. Cell. 183:28–45. doi:10.1016/j.cell.2020.09.014.

Moradi-Fard, S., J. Sarthi, M. tittel-Elmer, M. Lalonde, E. Cusanelli, P. Chartrand, and J.A. Cobb. 2016. Smc5/6 is a telomere-associated complex that regulates Sir4 binding and TPE. PLoS Genet. 12:e1006268. doi:10.1371/journal.pgen.1006268.

Murphy, S.E., and T.P. Levine. 2016. VAP, a Versatile Access Point for the Endoplasmic Reticulum: Review and analysis of FFAT-like motifs in the VAPome. Biochim. Biophys. Acta. 1861:952–961. doi:10.1016/j.bbalip.2016.02.009.

Ng, A.Q.E., A.Y.E. Ng, and D. Zhang. 2020. Plasma Membrane Furrows Control Plasticity of ER-PM Contacts. Cell Rep. 30:1434–1446. doi:10.1016/j.celrep.2019.12.098.

Ninova, M., K. Fejes Tóth, and A.A. Aravin. 2019. The control of gene expression and cell identity by H3K9 trimethylation. Development. 146:dev181180. doi:10.1242/dev.181180.

Panse, V.G., U. Hardeland, T. Werner, B. Kuster, and E. Hurt. 2004. A proteome-wide approach identifies sumoylated substrate proteins in yeast. J. Biol. Chem. 279:41346–41351. doi:10.1074/jbc.M407950200.

Poleshko, A., K.M. Mansfield, C.C. Burlingame, M.D. Andrake, N.R. Shah, and R.A. Katz. 2013. The human protein PRR14 tethers heterochromatin to the nuclear lamina during interphase and mitotic exit. Cell Rep. 5:292–301. doi:10.1016/j.celrep.2013.09.024.

Poleshko, A., C.L. Smith, S.C. Nguyen, P. Sivaramakrishnan, K.G. Wong, J.I. Murray, M. Lakadamyali, E.F. Joyce, R. Jain, and J.A. Epstein. 2019. H3K9me2 orchestrates inheritance of spatial positioning of peripheral heterochromatin through mitosis. eLife. 8:e49278. doi:10.7554/eLife.49278.

Politz, J.C., D. Scalzo, and M. Groudine. 2013. Something silent this way forms: the functional organization of the repressive nuclear compartment. Annu. Rev. Cell Dev. Biol. 29:241–270. doi:10.1146/annurev-cellbio-101512-122317.

Ptak, C., J.D. Aitchison, and R.W. Wozniak. 2014. The multifunctional nuclear pore complex: a platform for controlling gene expression. Curr. Opin. Cell Biol. 28:46–53. doi:10.1016/j.ceb.2014.02.001.

Ptak C., and R.W. Wozniak. 2016. Nucleoporins and chromatin metabolism. Curr. Opin. Cell Biol. 40: 153–160. doi:10.1016/j.ceb.2016.03.024.

Psakhye, I., and S. Jentsch. 2012. Protein group modification and synergy in the SUMO pathway as exemplified in DNA repair. Cell. 151: 807–820. doi:10.1016/j.cell.2012.10.021.

Rosonina, E., S.M. Duncan, and J.L. Manley. 2010. SUMO functions in constitutive transcription and during activation of inducible genes in yeast. Genes Dev. 24:1242–1252. doi:10.1101/gad.1917910.

Saik, N.O., N. Park, C. Ptak, N. Adames, J.D. Aitchison, and R.W. Wozniak. 2020. Recruitment of an Activated Gene to the Yeast Nuclear Pore Complex Requires Sumoylation. Front. Genet. 11:174. doi:10.3389/fgene.2020.00174.

Sikorski, R.S., and P. Hieter. 1989. A system of shuttle vectors and yeast host strains designed for efficient manipulation of DNA in Saccharomyces cerevisiae. Genetics. 122:19–27.

Smoyer, C.J., S.S. Katta, J.M. Gardner, L. Stoltz, S. McCroskey, W.D. Bradford, M. McClain, S.E. Smith, B.D. Slaughter, J.R. Unruh, and S.L. Jaspersen. 2016. Analysis of membrane proteins localizing to the inner nuclear envelope in living cells. J. Cell Biol. 215:575–590. doi:10.1083/jcb.201607043.

Stefan, C.J., A.G. Manford, D. Baird, J. Yamada-Hanff, Y. Mao, and S.D. Emr. 2011. Osh proteins regulate phosphoinositide metabolism at ER–plasma membrane contact sites. Cell. 144:389–401. doi:10.1016/j.cell.2010.12.034.

Taddei, A., F. Hediger, F.R. Neumann, C. Bauer, and S.M. Gasser. 2004. Separation of silencing from perinuclear anchoring functions in yeast Ku80, Sir4 and Esc1 proteins. EMBO J. 23:1301–1312. doi:10.1038/sj.emboj.7600144.

Taddei, A., and S.M. Gasser. 2012. Structure and function in the budding yeast nucleus. Genetics. 192:107–129. doi:10.1534/genetics.112.140608.

Texari, L., G. Dieppois, P. Vinciguerra, M.P. Contreras, A. Groner, A. Letourneau, and F. Stutz. 2013. The nuclear pore regulates GAL1 gene transcription by controlling the localization of the SUMO protease Ulp1. Mol. Cell. 51:807–818. doi:10.1016/j.molcel.2013.08.047.

Texari, L., and F. Stutz. 2015. Sumoylation and transcription regulation at nuclear pores. Chromosoma. 124:45–56. doi:10.1007/s00412-014-0481-x.

Van Damme, E., K. Laukens, T.H. Dang, and X. Van Ostade. 2010. A manually curated network of the PML nuclear body interactome reveals an important role for PML-NBs in SUMOylation dynamics. Int. J. Biol. Sci. 6:51–67. doi:10.7150/ijbs.6.51.

Van de Vosse, D.W., Y. Wan, D.L. Lapetina, W.M. Chen, J.H. Chiang, J.D. Aitchison, and R.W. Wozniak. 2013. A role for the nucleoporin Nup170p in chromatin structure and gene silencing. Cell. 152:969–983. doi:10.1016/j.cell.2013.01.049.

Weber-Boyvat, M., H. Kentala, J. Lilja, T. Vihervaara, R. Hanninen, Y. Zhou, J. Peranen, T.A. Nyman, J. Ivaska, and V.M. Olkkonen. 2015. OSBP-related protein 3 (ORP3) coupling with VAMP associated protein A regulates R-Ras activity. Exp. Cell Res. 331:278–291. doi:10.1016/j.yexcr.2014.10.019.

Wohlschlegel, J.A., E.S. Johnson, S.I. Reed, and J.R. 3rd Yates. 2004. Global analysis of protein sumoylation in Saccharomyces cerevisiae. J. Biol. Chem. 279:45662–45668. doi:10.1074/jbc.M409203200.

Wurzenberger, C., and D.W. Gerlich. 2011. Phosphatases: providing safe passage through mitotic exit. Nat. Rev. Mol. Cell Biol. 12:469–482. doi:10.1038/nrm3149.

Wykoff, D.D., and E.K. O’Shea. 2005. Identification of sumoylated proteins by systematic immunoprecipitation of the budding yeast proteome. Mol. Cell. Proteomics. 4:73–83. doi:10.1074/mcp.M400166-MCP200.

Zhao, Q., X. Y. Xie, Y. Zheng, S. Jiang, W. Liu, W. Mu, Z. Liu, Y. Zhao, Y. Xue, and J. Ren. 2014. GPS-SUMO: a tool for the prediction of sumoylation sites and SUMO-interaction motifs. Nucleic Acids Res. 42:(Web Server issue) W325–W330. doi:10.1093/nar/gku383.

Zhou, W., J.J. Ryan, and H. Zhou. 2004. Global analyses of sumoylated proteins in Saccharomyces cerevisiae. Induction of protein sumoylation by cellular stresses. J. Biol. Chem. 279:32262–32268. doi:10.1074/jbc.M404173200.

