## Supplementary material for "Phosphorylation-dependent mitotic SUMOylation drives nuclear envelope-chromatin interactions": All supplementary material

#### Supplemental Figure Legends

**Figure S1. Siz2 localization and post-translational modification.** (**A** and **G**) α-factor arrest-release assays were carried out as in Fig. 1 using *siz1Δ* cells (**A**), as well as Siz2-V$5_3$, siz$2^{S527A}$-V$5_3$, and siz$2^{S674A}$-V$5_3$ producing cells (**G)**. Cell lysates were analyzed by western blotting using anti-SUMO, Clb2, and Gsp1 (load control) antibodies, as well as a V5 antibody in **G**, as indicated. (**B**) Cell lysates derived from asynchronous cultures of WT, *siz1Δ, siz2Δ* , and *mms21$^{1-184}$-V5$_3$* (Mms21 derivative deficient in SUMO E3 ligase activity) cells were assessed by western blotting using an anti-SUMO antibody to assess SUMO conjugate profiles. Gsp1 is a loading control. In (**A**, **B** and **G**) asterisks identify expected positions of mitotic, Scs2-Siz2 complex-dependent SUMO conjugates. In **G**, dots indicate the expected position of mitotically phosphorylated Siz2-V$5_3$. Molecular mass markers are shown in kDa. (**C**) Epifluorescence images of *siz1Δ* and *mms21$^{1-184}$-V5$_3$* cells were analyzed by immunofluorescence using anti-SUMO antibodies (SUMO) as described in Fig.1. DAPI staining identified nuclear position. (**D**) Shown are epifluorescence images of representative G1/S- and M-phase WT cells producing GFP-Siz2 and nucleolar Nop56-mCherry. Merged images show that GFP-Siz2 is largely excluded from the nucleolus. (**E**) GFP-Siz2 intensity (y-axis) through the nucleus (x-axis) was assessed by line scan in a representative G1- and M-phase cell using rendered images shown in Fig. 2 C. The line through each nucleus identifies the path of the scan. (**F**) WT mitotic cell lysates were isolated at 60 min post-release from α-factor arrest (see Fig. 1 B). Proteins were extracted and solubilized in buffer lacking (-PPase) or containing protein phosphatase (+PPase). Samples were analyzed by western blotting using anti-V5 and Gsp1 (loading control) antibodies. Mobility of modified Siz2-V$5_3$ is increased by phosphatase treatment. (**G**) Epifluorescence images of representative mitotic cells producing GFP-Siz2, GFP- siz$2^{S527A}$ and GFP-siz$2^{S674A}$ along with the NE/ER marker Sur4-mCherry. Bar, 2 μm.

**Figure S2. Siz2 directs mitosis specific SUMOylation of the VAP family member Scs2.** (**A**) Shown are epifluorescence images of a mitotic cell producing GFP-Siz2 and scs$2^{1-225}$-mCherry. (**B**), The position, sequence, and score (Murphy and Levine, 2016) of putative Siz2 FFAT-like motifs are shown. An optimal FFAT sequence is shown for comparison. (**C** and **E**) Using the split-superfolder GFP system (see Fig. 3 A), the INM association of the *scs2* MSP (**C**) domain mutant and the *scs2* SUMO site mutant (d) were assessed in cells producing GFP$_{1-10}$-

scs2$^{K84D/L86D}$ or GFP$_{1-10}$-scs2$^{K180R}$ and the plasmid-encoded GFP$_{11}$-mCherry-Pus1 reporter. Bar, 2 μm. (**D**) α-factor arrest-release assays were carried out as in Fig. 1 using *ulp1*$^{K352E/Y583H}$*-V5*$_3$ mutant stains producing Siz2-V5$_3$, scs2$^{K84D/L86D}$, or siz2$^{A569D}$-V5$_3$ as indicated. Cell lysates were analyzed by western blotting using anti-V5, SUMO, Clb2, and Gsp1 (load control) antibodies. Asterisks identify expected positions of mitotic, Scs2-Siz2 complex-dependent SUMO conjugates. Molecular mass markers are shown in kDa.

**Figure S3. INM localized Scs2-Siz2 is required for telomere tethering during mitosis and G1-phase of the cell cycle.** (**A-C**), NE tethering of Tel16R in WT and *siz2*$^{S522A}$*-V5*$_3$ cells (**A**), as well as was NE tethering of Tel14L in WT (**B**, **C**) *ulp1*$^{K352E/Y583H}$*-V5*$_3$, *ulp1*$^{K352E/Y583H}$*-V5*$_3$ *siz2*$^{S522A}$*-V5*$_3$ (**B**) *yku70Δ*, *yku80Δ*, and *sir4Δ* cells (**C**) were examined as described in Fig. 5. (**D**) Shown are epifluorescence images of WT cells producing Sir4-GFP and NE/ER localized Sur4-mCherry. Merged images were used to assess the relative position of Sir4-GFP foci with respect to the NE (identified by Sur4-mCherry). Images were rendered using the unsharp mask filter in Image J. Bar, 2 μm. Bar graphs show the percentage of total Sir4-GFP foci at the NE (middle panel) and the average number of Sir4-GFP foci per nucleus (right panel) in WT and *siz2*Δ cells at the indicated cell cycle stage. Note, cells in anaphase/telophase show double the number of Sir4-GFP foci as seen in G1- and S-phase cells. Graphs in (**A-D**) represent data from at least three biological replicates. n = 50 cells/replicate/cell cycle stage. Error bars – SD. Asterisks – significant difference relative to WT using a two-tailed student's t-test. *$p \leq 0.05$, **$p \leq 0.01$, ***$p \leq 0.001$. **e**, Sir4-PrA binding to chromatin adjacent to Tel6R was assessed by ChIP and qRT-PCR analysis using asynchronous cultures of WT, *ulp1*$^{K352E/Y583H}$*-V5*$_3$, *siz2*$^{S522A}$*-V5*$_3$, and *ulp1*$^{K352E/Y583H}$*-V5*$_3$ *siz2*$^{S522A}$*-V5*$_3$, cells. Graphs represent at least three biological replicates. Error bars - SEM. Asterisks - significant change relative to WT using a two-tailed student's t-test. *$p \leq 0.05$, **$p \leq 0.01$.

**Figure S4. The *sir4*$^{K1037R}$ mutation affects telomere tethering but not Sir4 NE localization.** (**A** and **D**) SUMO conjugates (**A**)**,** Sir4-V5$_3$, or sir4$^{K1037R}$-V5$_3$ (**E**) bound to chromatin adjacent to Tel6R was assessed by ChIP and qRT-PCR analysis using antibodies directed against SUMO or the V5 epitope and asynchronous cultures of the indicated strains. Graphs represent at least three biological replicates. Error bars - SEM. Asterisks - significant change relative to WT using a two-tailed student's t-test. (**B**) Tethering of Tel16R in WT and *sir4*$^{K1037R}$*-V5*$_3$ cells was examined as described in Fig. 5 A. (**C**) NE localization of Sir4-GFP and sir4$^{K1037R}$-GFP was assessed as described in Extended Data Fig. 4 C. Bar graphs show the percentage of total GFP foci at the NE (left panel) and the average number of foci per nucleus (right panel) at the indicated cell cycle stage. Cells in anaphase/telophase show double the number of Sir4-GFP foci as seen in G1- and S-phase cells. Note, WT data shown is the same as that in Extended Data Fig. 4 C**.** Graphs in (**B**) and (**C**) represent data from at least three biological replicates. n = 50 cells/replicate/cell cycle stage. Error bars – SD. Asterisks – significance significant difference relative to WT using a two-tailed student's t-test. *$p \leq 0.05$, **$p \leq 0.01$, ***$p \leq 0.001$.

**Figure S5. SUMOylation contributes to Sir4 subtelomeric chromatin association during mitosis.** To assess cell cycle stage of cultures used for cell-cycle dependent ChIP analysis (see Fig. 6 E), samples at each time point were analyzed by FACS to determine DNA content of cells in the population. The positions of 1n and 2n DNA peaks are shown. To the right, proteins derived from cell lysates harvested at the various time points were analyzed by western blotting using anti-V5, SUMO, Clb2, and Gsp1 (load control) antibodies. Asterisks identify expected positions of mitotic, Scs2-Siz2 dependent SUMO conjugates. Molecular mass markers are shown in kDa.

**Video 1. Dynamics of GFP-Siz2 localization throughout the cell cycle.** Cell-cycle dependent changes in Siz2 localization were visualized using time-lapse epifluorescence microscopy on cells producing GFP-Siz2. Images were acquired at 1.5 min intervals over a 30 min period. Highlighted are cells progressing through G1- and S-phase (yellow dots), entering mitosis (red dots), and exiting mitosis (green dots). Images were rendered using the unsharp mask filter in Image J and saved as a QuickTime movie.

Figure S1

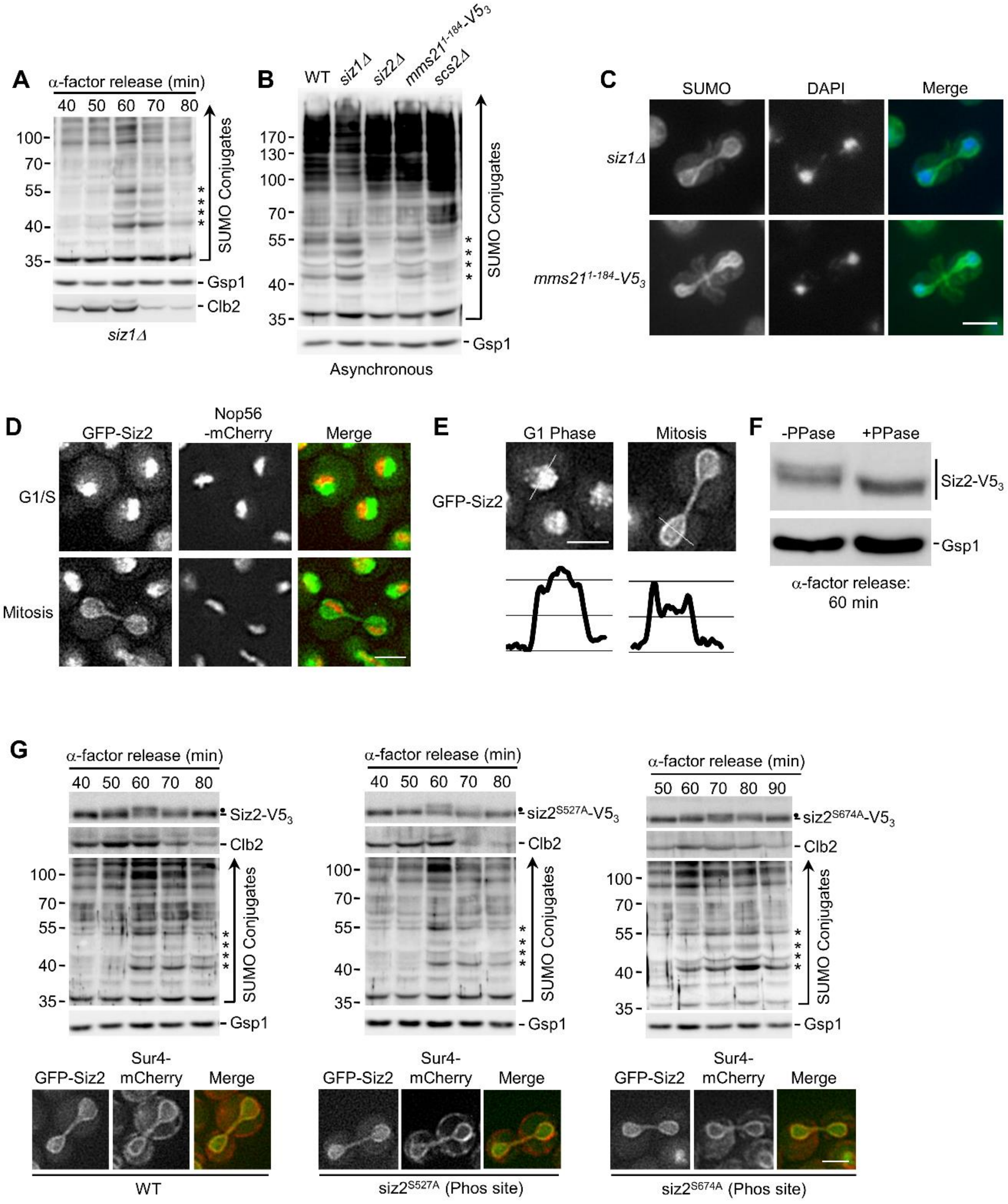
A
α-factor release (min)
40 50 60 70 80
100
70
55
40
35
SUMO Conjugates
Gsp1
Clb2
siz1Δ
B
WT siz1Δ siz2Δ mms21^1-184-V5_3 scs2Δ
170
130
100
70
55
40
35
SUMO Conjugates
Gsp1
Asynchronous
C
SUMO DAPI Merge
siz1Δ
mms21^1-184-V5_3
D
GFP-Siz2
Nop56 -mCherry
Merge
G1/S
Mitosis
E
G1 Phase
Mitosis
GFP-Siz2
F
-PPase +PPase
Siz2-V5_3
Gsp1
α-factor release: 60 min
G
α-factor release (min)
40 50 60 70 80
Siz2-V5_3
Clb2
100
70
55
40
35
SUMO Conjugates
Gsp1
α-factor release (min)
40 50 60 70 80
siz2^S527A-V5_3
Clb2
SUMO Conjugates
Gsp1
α-factor release (min)
50 60 70 80 90
siz2^S674A-V5_3
Clb2
SUMO Conjugates
Gsp1
GFP-Siz2 Sur4-mCherry Merge
WT
siz2^S527A (Phos site)
siz2^S674A (Phos site)

### Figure S2

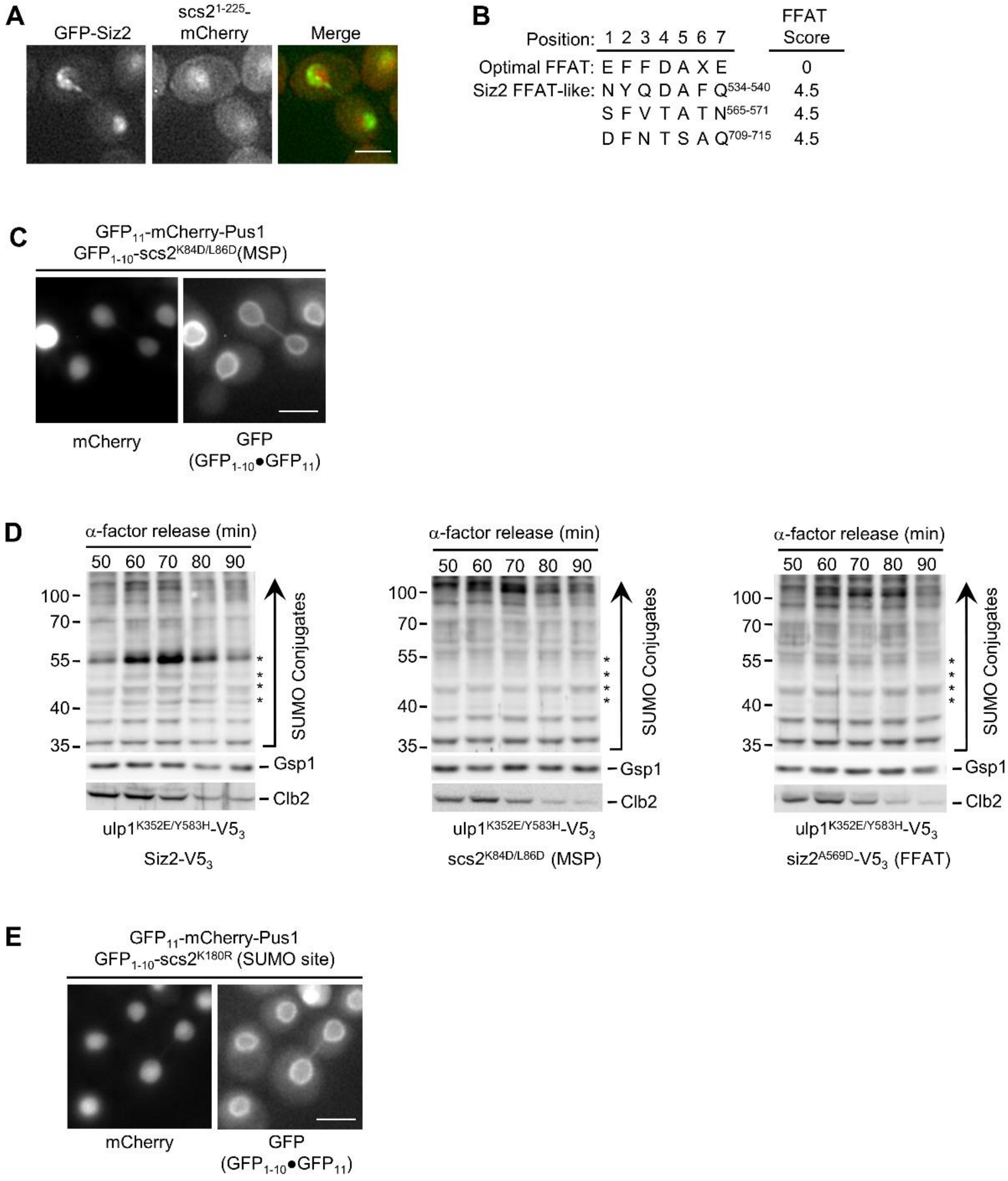
A
GFP-Siz2
scs2^1-225-mCherry
Merge
B
Position: 1 2 3 4 5 6 7
FFAT Score
Optimal FFAT: E F F D A X E 0
Siz2 FFAT-like: N Y Q D A F Q534-540 4.5
S F V T A T N565-571 4.5
D F N T S A Q709-715 4.5
C
GFP11-mCherry-Pus1
GFP1-10-scs2K84D/L86D(MSP)
mCherry
GFP (GFP1-10•GFP11)
D
α-factor release (min)
50 60 70 80 90
100
70
55
40
35
SUMO Conjugates
Gsp1
Clb2
ulp1K352E/Y583H-V53
Siz2-V53
scs2K84D/L86D (MSP)
siz2A569D-V53 (FFAT)
E
GFP11-mCherry-Pus1
GFP1-10-scs2K180R (SUMO site)
mCherry
GFP (GFP1-10•GFP11)

### Figure S3

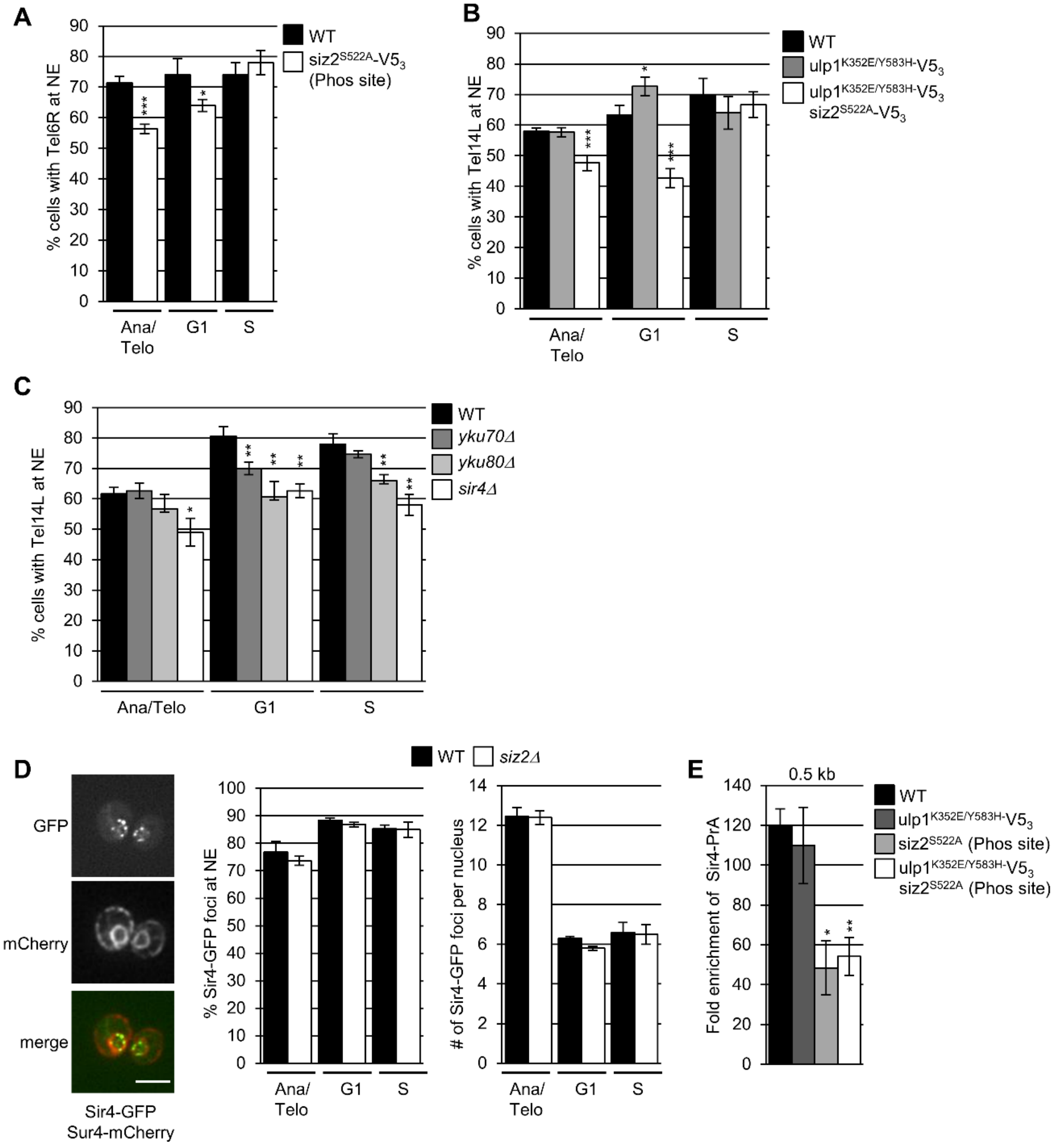

A
% cells with Tel6R at NE
WT
siz2S522A-V53 (Phos site)
Ana/ Telo
G1
S
B
% cells with Tel14L at NE
WT
ulp1K352E/Y583H-V53
ulp1K352E/Y583H-V53 siz2S522A-V53
C
% cells with Tel14L at NE
WT
yku70Δ
yku80Δ
sir4Δ
Ana/Telo
D
GFP
mCherry
merge
Sir4-GFP Sur4-mCherry
WT
siz2Δ
% Sir4-GFP foci at NE
### of Sir4-GFP foci per nucleus
E
0.5 kb
Fold enrichment of Sir4-PrA
WT
ulp1K352E/Y583H-V53
siz2S522A (Phos site)
ulp1K352E/Y583H-V53 siz2S522A (Phos site)

Figure S4

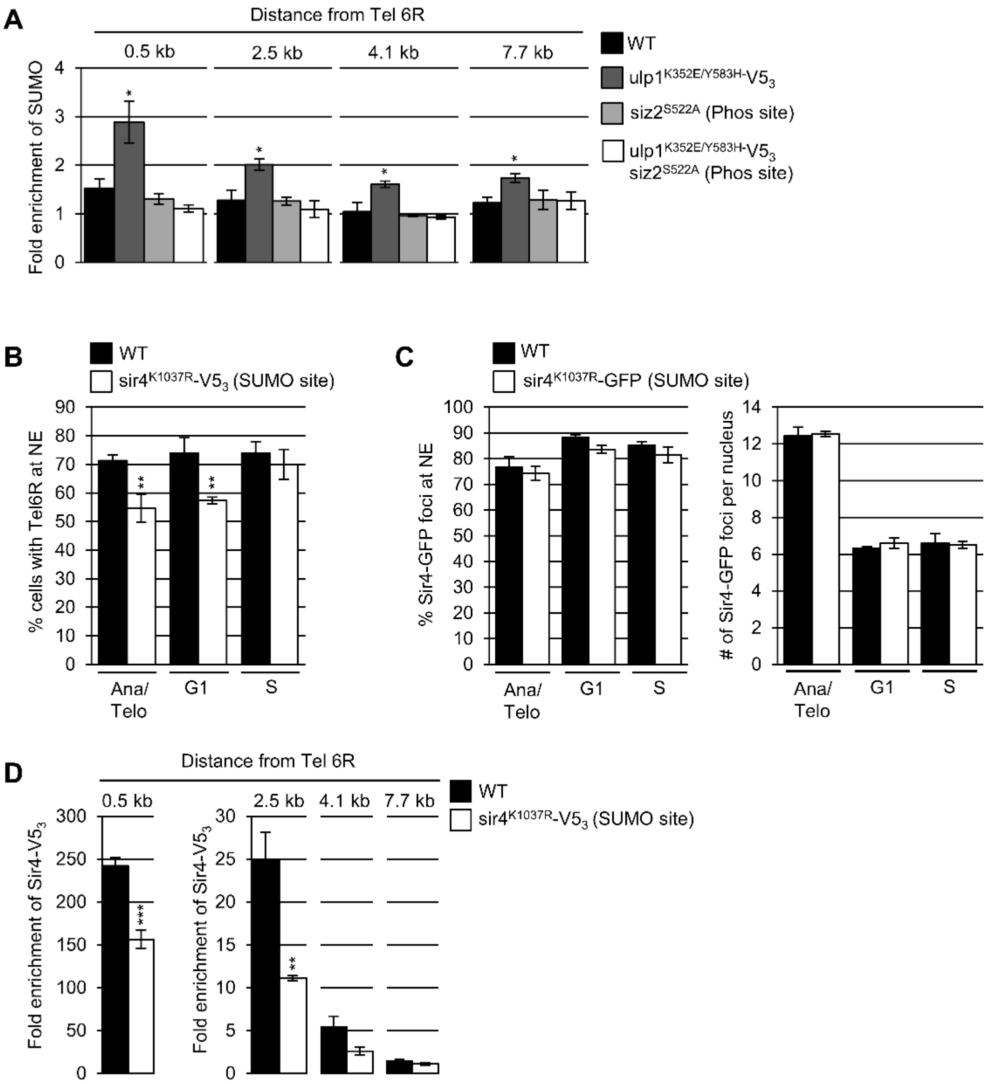
A
Distance from Tel 6R
0.5 kb
2.5 kb
4.1 kb
7.7 kb
Fold enrichment of SUMO
WT
ulp1K352E/Y583H-V53
siz2S522A (Phos site)
ulp1K352E/Y583H-V53 siz2S522A (Phos site)
B
WT
sir4K1037R-V53 (SUMO site)
% cells with Tel6R at NE
Ana/ Telo
G1
S
C
WT
sir4K1037R-GFP (SUMO site)
% Sir4-GFP foci at NE
### of Sir4-GFP foci per nucleus
D
Distance from Tel 6R
WT
sir4K1037R-V53 (SUMO site)
Fold enrichment of Sir4-V53

### Figure S5

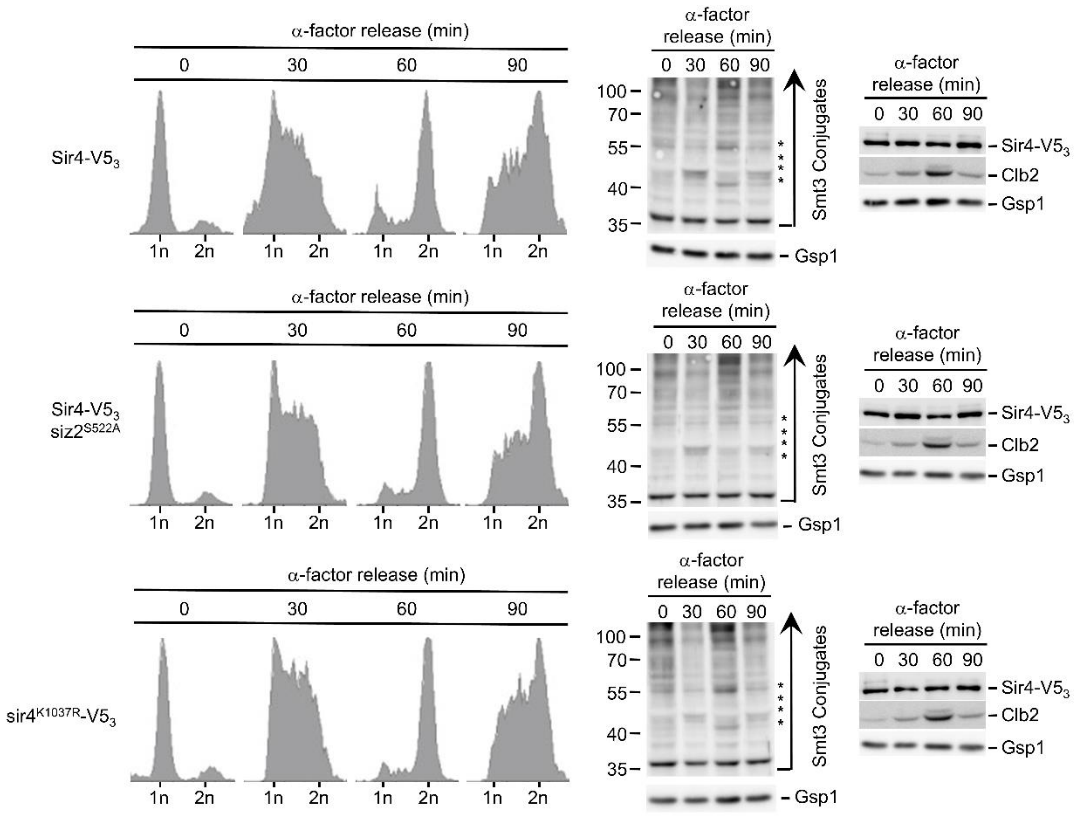
α-factor release (min)
0
30
60
90
Sir4-V5$_3$
1n 2n
α-factor release (min)
0 30 60 90
100
70
55
40
35
Smt3 Conjugates
Gsp1
α-factor release (min)
0 30 60 90
Sir4-V5$_3$
Clb2
Gsp1
Sir4-V5$_3$ siz2$^{S522A}$
sir4$^{K1037R}$-V5$_3$

### Table S1. Strains List

| | Relevant Genotype | Source |
|---|---|---|
| BY4741* | *MAT**a** his3Δ1 leu2Δ0 ura3Δ0 met15Δ0* | Brachmann et al., 1998 |
| NS2001 | *MAT**a** bar1Δ::NAT* | This study |
| NS2099 | *MAT**a** SIZ2-V5$_3$-HIS scs2Δ::KAN bar1Δ::NAT* | This study |
| CPY4082 | *MAT**a** SIZ2-V5$_3$-HIS KAN-scs2$^{K180R}$ bar1Δ::NAT* | This study |
| CPY3908 | *MAT**a** ulp1$^{K352E}$-KAN bar1Δ::NAT* | This study |
| CPY3847 | *MAT**a** ulp1$^{K352E/Y583H}$-V53-HIS bar1Δ::NAT* | This study |
| CPY3864 | *MAT**a** KAN-SCS2pr-HA$_3$-SCS2 ulp1$^{K352E/Y582H}$-V5$_3$-HIS bar1Δ::NAT* | This study |
| CPY4163 | *MAT**a** SCS2-V5$_3$-KAN ulp1$^{K352E/Y583H}$-V5$_3$-HIS bar1Δ::NAT* | This study |
| CPY4006 | *MAT**a** scs2Δ::NAT* | This study |
| CPY3888 | *MAT**a** KAN-scs2$^{K180R}$* | This study |
| NS2018 | *MAT**a** siz2Δ::KAN bar1Δ::NAT* | This study |
| CPY4004 | *MAT**a** siz2Δ::KAN* | This study |
| CPY3784 | *MAT**a** HIS-SIZ2pr-GFP-SIZ2 SUR4-mCherry-NAT* | This study |
| CPY3909 | *MAT**a** SIZ2-V5$_3$-KAN bar1Δ::NAT* | This study |
| CPY3814 | *MAT**a** siz2$^{S522A}$-V5$_3$-KAN bar1Δ::NAT* | This study |
| CPY3867 | *MAT**a** HIS-SIZ2pr-GFP-siz2 $^{S522A}$-HPH SUR4-mCherry-NAT* | This study |
| CPY4325 | *MAT**a** siz1Δ::KAN bar1Δ::NAT* | This study |
| CPY3134 | *MAT**a** siz1Δ::KAN* | This study |
| CPY4193 | *MATα mms211-184-V53-KAN* | This study |
| CPY3801 | *MAT**a** HIS- SIZ2pr-GFP-SIZ2 NOP56-mCherry-NAT* | This study |
| CPY3851 | *MAT**a** siz2$^{S527A}$-V5$_3$-KAN bar1Δ::NAT* | This study |
| CPY3869 | *MAT**a** HIS- SIZ2pr-GFP-siz2 $^{S527A}$-HPH SUR4-mCherry-NAT* | This study |
| CPY3641 | *MAT**a** siz2$^{S674A-}$V5$_3$-HIS bar1Δ::NAT* | This study |
| CPY3841 | *MAT**a** HIS-SIZ2pr-GFP-siz2 $^{S674A}$-HPH SUR4-mCherry-NAT* | This study |
| CPY4100 | *MAT**a** NAT-CDC42pr-GFP$_{1-10}$-SCS2 pRS315-GFP$_{11}$-mCherry-Pus1* | This study |
| CPY4101 | *MAT**a** NAT-CDC42pr-GFP$_{1-10}$-SCS2 pRS315-GFP$_{11}$-mCherry-Hxk1* | This study |
| NS2911 | *MAT**a** SCS2-TAP-HIS SIZ2-V5$_3$-KAN* | This study |
| NS2917 | *MAT**a** SCS2-TAP-HIS siz2$^{S522A}$-V5$_3$-KAN* | This study |
| CPY3952 | *MAT**a** HIS- SIZ2pr-GFP-SIZ2 SUR4-mCherry-NAT scs2Δ::KAN* | This study |
| CPY4061 | *MAT**a** SIZ2-V5$_3$-HIS KAN-scs2$^{K84D/L86D}$ bar1Δ::NAT* | This study |
| CPY4036 | *MAT**a** HIS-SIZ2pr-GFP-SIZ2 SUR4-mCherry-NAT KAN-scs2$^{K84D/L86D}$* | This study |
| CPY4066 | *MAT**a** siz2$^{A569D}$-V5$_3$-KAN bar1Δ::NAT* | This study |
| CPY4089 | *MAT**a** HIS-SIZ2pr-GFP- siz2$^{A569D}$-HPH SUR4-mCherry-NAT* | This study |
| CPY4028 | *MAT**a** HIS-SIZ2pr-GFP-SIZ2 scs2$^{1-225}$-mCherry-NAT* | This study |
| CPY4225 | *MAT**a** NAT-CDC42pr-GFP$_{1-10}$-scs2$^{K84D/L86D}$ pRS315-GFP$_{11}$-mCherry-Pus1* | This study |
| CPY3911 | *MAT**a** SIZ2-V5$_3$-KAN ulp1$^{K352E/Y583H}$-V53-HPH bar1Δ::NAT* | This study |
| CPY4030 | *MAT**a** KAN-scs2$^{K84D/L86D}$ ulp1$^{K352E/Y583H}$-V5$_3$-HIS bar1Δ::NAT* | This study |
| CPY4072 | *MAT**a** siz2$^{A569D}$-V5$_3$-KAN ulp1$^{K352E/Y583H}$-V5$_3$-HIS bar1Δ::NAT* | This study |
| CPY3894 | *MAT**a** HIS-SIZ2pr-GFP-SIZ2-HPH SUR4-mCherry-NAT KAN-scs2$^{K180R}$* | This study |
| CPY3915 | *MAT**a** siz2$^{I472/473A}$-V5$_3$-KAN bar1Δ::NAT* | This study |
| CPY3835 | *MAT**a** HIS-SIZ2pr-GFP-siz2$^{I472/473A}$-HPH SUR4-mCherry-NAT* | This study |
| CPY4092 | *MAT**a** siz2$^{V720/721A}$-V5$_3$-KAN bar1Δ::NAT* | This study |
| CPY3837 | *MAT**a** HIS-SIZ2pr-GFP-siz2$^{V720/721A}$-HPH SUR4-mCherry-NAT* | This study |
| CPY3917 | *MAT**a** siz2$^{I472/473A}$-V5$_3$-KAN ulp1$^{K352E/Y583H}$-V5$_3$-HPH bar1Δ::NAT* | This study |
| CPY3835 | *MAT**a** HIS-SIZ2pr-GFP-siz2$^{I472/473A}$-HPH SUR4-mCherry-NAT ulp1$^{K352E/Y583H}$-V5$_3$-KAN* | This study |
| CPY4221 | *MAT**a** NAT-CDC42pr-GFP$_{1-10}$-scs2$^{K180R}$ pRS315-GFP$_{11}$-mCherry-Pus1* | This study |
| CPY3856 | *MAT**a** HIS-SIZ2pr-GFP-siz2 $^{I472/473A, V720/721A}$-HPH SUR4-mCherry-NAT* | This study |
| DVY1534 | *MAT**a** W303 leu2-3,112 ura3-1 trp1-1 ade2-1 can1-100 TelXIV-L::256xlacOR-TRP1 HIS3::GFP-lacI-HIS3 SEC63-GFP-NAT* | Van de Vosse et al., 2013 |
| CPY3981 | *MAT**a** W303 leu2-3,112 ura3-1 trp1-1 ade2-1 can1-100 TelXIV-L::256xlacO-TRP1 HIS3::GFP-lacI-HIS3 SEC63-GFP-NAT siz2$^{S522A}$-V5$_3$-KAN* | This study |
| CPY4070 | *MAT**a** W303 leu2-3,112 ura3-1 trp1-1 ade2-1 can1-100 TelXIV-L::256xlacO-TRP1 HIS3::GFP-lacI-HIS3 SEC63-GFP-NAT siz2$^{A569D}$-V5$_3$-KAN* | This study |
| CPY3776 | *MAT**a** W303 leu2-3,112 ura3-1 trp1-1 ade2-1 can1-100 TelXIV-L::256xlacO-TRP1 HIS3::GFP-lacI-HIS3 SEC63-GFP-NAT siz2$^{I472/473A}$-V5$_3$-KAN* | This study |

| | | |
|---|---|---|
| CPY3992 | *MAT**a** W303 leu2-3,112 ura3-1 trp1-1 ade2-1 can1-100 TelXIV-L::256xlacO-TRP1 HIS3::GFP-lacI-HIS3 SEC63-GFP-NAT KAN-scs2*$^{K84D/L86D}$ | This study |
| NS2418 | *MAT**a** W303 leu2-3,112 ura3-1 trp1-1 ade2-1 can1-100 TelXIV-L::256xlacO-TRP1 HIS3::GFP-lacI-HIS3 SEC63-GFP-NAT KAN-scs2*$^{K180R}$ | This study |
| CPY4049 | *MAT**a** SIR4-V5*$_3$*-KAN* | This study |
| NS2500 | *MAT**a** SIR4-V5*$_3$*-KAN siz2*$^{S522A}$*-NAT* | This study |
| NS2506 | *MAT**a** SIR4-V5*$^3$*-KAN NAT-scs2*$^{K84D/L86D}$ | This study |
| NS2504 | *MAT**a** SIR4-V5*$^3$*-KAN NAT-scs2*$^{K180R}$ | This study |
| NS3522 | *MAT**a** W303 leu2-3,112 ura3-1 trp1-1 ade2-1 can1-100 TelXIV-L::256xlacO-TRP1 HIS3::GFP-lacI-HIS3 SEC63-GFP-NAT ulp1*$^{K352E/Y583H}$*-V5*$_3$*-HPH* | This study |
| NS3524 | *MAT**a** W303 leu2-3,112 ura3-1 trp1-1 ade2-1 can1-100 TelXIV-L::256xlacO-TRP1 HIS3::GFP-lacI-HIS3 SEC63-GFP-NAT ulp1*$^{K352E/Y583H}$*-V5*$_3$*-HPH siz2*$^{S522A}$*-V5*$_3$*-KAN* | This study |
| NS3206 | *MAT**a** W303 leu2-3,112 ura3-1 trp1-1 ade2-1 can1-100 TelXIV-L::256xlacO-TRP1 HIS3::GFP-lacI-HIS3 SEC63-GFP-NAT yku80Δ::KAN* | This study |
| DVY1539.1 | *MAT**a** W303 leu2-3,112 ura3-1 trp1-1 ade2-1 can1-100 TelXIV-L::256xlacO-TRP1 HIS3::GFP-lacI-HIS3 SEC63-GFP-NAT yku70Δ::KAN* | Van de Vosse et al., 2013 |
| DVY1539 | *MAT**a** W303 leu2-3,112 ura3-1 trp1-1 ade2-1 can1-100 TelXIV-L::256xlacO-TRP1 HIS3::GFP-lacI-HIS3 SEC63-GFP-NAT sir4Δ::KAN* | Van de Vosse et al., 2013 |
| NS2078 | *MAT**a** SIR4-eGFP-HIS SUR4-mCherry-NAT* | This study |
| NS2144 | *MAT**a** SIR4-eGFP-HIS SUR4-mCherry-NAT siz2Δ::KAN* | This study |
| DVY2055 | *MAT**a** W303 leu2-3,112 ura3-1 trp1-1 ade2-1 can1-100 TelVI-R::256xlacO-TRP1 HIS3::GFP-lacI-HIS3 SEC63-GFP-NAT* | Van de Vosse et al., 2013 |
| CPY3988 | *MAT**a** W303 leu2-3,112 ura3-1 trp1-1 ade2-1 can1-100 TelVI-R::256xlacO-TRP1 HIS3::GFP-lacI-HIS3 SEC63-GFP-NAT siz2*$^{S522A}$*-V5*$_3$*-KAN* | This study |
| NS2111 | *MAT**a** SIR4-PrA-HIS* | This study |
| NS3447 | *MAT**a** SIR4-PrA-HIS ulp1*$^{K352E/Y583H}$*-V5*$_3$*-KAN* | This study |
| NS3513 | *MAT**a** SIR4-PrA-HIS siz2*$^{S522A}$*-NAT* | This study |
| NS3511 | *MAT**a** SIR4-PrA-HIS ulp1*$^{K352E/Y583H}$*-V5*$_3$*-KAN siz2*$^{S522A}$*-NAT* | This study |
| CPY4050 | *MAT**a** SIR4-V5*$_3$*-KAN SMT3pr-HIS*$_8$*-SMT3-HPH* | This study |
| CPY4053 | *MAT**a** SIR4-V5*$_3$*-KAN SMT3pr-HIS*$_8$*-SMT3-HPH siz2S*$^{522A}$*-NAT* | This study |
| CPY4056 | *MAT**a** SIR4-V5*$_3$*-KAN SMT3pr-HIS*$_8$*-SMT3-HPH NAT-scs2*$^{K180R}$ | This study |
| CPY4057 | *MAT**a** SIR4-V5*$_3$*-KAN SMT3pr-HIS*$_8$*-SMT3-HPH NAT-scs2*$^{K84D/L86D}$ | This study |
| NS3501 | *MAT**a** SIR4-V5*$_3$*-KAN SMT3pr-HIS*$_8$*-SMT3-HPH ulp1*$^{K352E/Y583H}$*-V5*$_3$*-HIS* | This study |
| NS3503 | *MAT**a** SIR4-V5*$_3$*-KAN SMT3pr-HIS*$_8$*-SMT3-HPH ulp1*$^{K352E/Y583H}$*-V5*$_3$*-HIS siz2*$^{S522A}$*-NAT* | This study |
| CPY4051 | *MAT**a** sir4*$^{K1037R}$*-V5*$_3$*-KAN SMT3pr-HIS*$_8$*-SMT3-HPH* | This study |
| CPY4008 | *MAT**a** W303 leu2-3,112 ura3-1 trp1-1 ade2-1 can1-100 TelXIV-L::256xlacO-TRP1 HIS3::GFP-lacI-HIS3 SEC63-GFP-NAT sir4*$^{K1037R}$*-V5*$_3$*-KAN* | This study |
| CPY4014 | *MAT**a** sir4*$^{K1037R}$*-V5*$_3$*-KAN* | This study |
| NS2705 | *MAT**a** SIR4-V5*$_3$*-KAN bar1Δ::NAT* | This study |
| NS2705 | *MAT**a** SIR4-V5*$_3$*-KAN siz2*$^{S522A}$*-NAT bar1Δ::NAT* | This study |
| CPY4244 | *MAT**a** W303 leu2-3,112 ura3-1 trp1-1 ade2-1 can1-100 TelVI-R::256xlacO-TRP1 HIS3::GFP-lacI-HIS3 SEC63-GFP-NAT sir4*$^{K1037R}$*-V5*$_3$*-KAN* | This study |
| NS2633 | *MAT**a** sir4*$^{K1037R}$*-eGFP-HIS SUR4-mCherry-NAT* | This study |
| NS2709 | *MAT**a** sir4*$^{K1037R}$*-V5*$_3$*-KAN bar1Δ::NAT* | This study |
| NS2433 | *MAT**a** SIR4-V5*$_3$*-KAN siz2Δ::HPH* | This study |
| NS2496 | *MAT**a** SIR4-V5*$_3$*-KAN scs2Δ::NAT* | This study |

*unless otherwise indicated all strains were derived from BY4741 and BY4742 with strains derived from dissections not genotyped for met15Δ0 or lys2Δ0.
